# Postnatal lymph node expansion of stromal progenitors towards reticular and CD34^+^ stromal cell subsets is determined by distinct transcriptional programs

**DOI:** 10.1101/2021.06.06.447189

**Authors:** Joern Pezoldt, Carolin Wiechers, Maria Litovchenko, Marjan Biocanin, Mangge Zou, Katarzyna Sitnik, Michael Beckstette, Wanze Chen, Vincent Gardeux, Stefan Floess, Maria Ebel, Julie Russeil, Panagiota Arampatzi, Ehsan Vafardanejad, Antoine-Emmanuel Saliba, Bart Deplancke, Jochen Huehn

## Abstract

Gut-draining mesenteric lymph nodes (mLN) provide the framework and microenvironment to shape intestinal adaptive immune responses. We previously delineated transcriptional signatures in LN stromal cells (SC), pointing to tissue-specific variability in composition and immuno-modulatory function of SCs.

Here, we dissect the tissue-specific epigenomic DNA accessibility and CpG methylation landscape of LN non-endothelial SCs and identify a microbiota-independent core epigenomic signature of LN SCs. By combined analysis of transcription factor (TF) binding sites together with the gene expression profiles of non-endothelial SCs, we delineated TFs poising skin-draining peripheral LN (pLN) SCs for pro-inflammatory responses. Furthermore, using scRNA-seq, we dissected the developmental trajectory of mLN SCs derived from postnatal to aged mice, identifying two distinct putative progenitors, namely CD34^+^ SC and fibroblastic reticular stromal cell (FRC) progenitors, which both feed the rapid postnatal LN expansion. Finally, we identified *Irf3* as a key differentiation TF inferred from the epigenomic signature of mLN SCs that is dynamically expressed along the differentiation trajectories of FRCs, and validated *Irf3* as a regulator of Cxcl9^+^ FRC differentiation.

Together, our data constitute a comprehensive transcriptional and epigenomic map of mLN development and dissect location-specific, microbiota-independent properties of mLN non-endothelial SCs. As such, our findings represent a valuable resource to identify core transcriptional regulators that impinge on the developing mLN early in life, thereby shaping long-lasting intestinal adaptive immune responses.

## Introduction

The mammalian immune system is tasked to detect pathogenic incursions and maintain balanced immune responses. Failure to achieve equilibrium can result in the development of local and systemic overreactions to self and foreign antigens or higher susceptibility to infections. As lymph nodes (LNs) are the initial hub translating early innate responses into lasting adaptive antigen-specific immunity, their tissue-specific modulation of developing immune responses is of essence to calibrate immune responses throughout life. Nonetheless, it is astounding that tissue-specific immune responses evolve in LNs given that they derive from seemingly identical postnatal LN anlagen that are positioned throughout the body.

LNs start to develop prenatally as early as embryonic day (E)13 in mice, in a process tightly regulated by mesenchymal lymphoid tissue organizer (LTo) and hematopoietic lymphoid tissue inducer (LTi) cells, yielding primordial LN anlagen.^1,2^ Initial recruitment and retention of LTis is mediated by endothelial LTos (eLTo).^3^ The same eLTos subsequently activate Cxcl13^+^ mesenchymal LTos (mLTo), further recruiting additional LTis in a Cxcl13^+^-driven manner.^2–4^ With the LN anlagen established by E17, containing a dense network of LTis and LTos,^5^ endothelial and mesenchymal cells further proliferate. Already at E18, lymphatic endothelial cells (LEC) have sufficiently expanded to envelope the core parenchyma of the LN, establishing the border to the surrounding tissue beneath the forming LN capsule.^6^ After birth, the influx of T cells and B cells drastically increases, requiring and driving the rapid expansion of the LN stromal cell (LNSC) compartment including non-endothelial SCs. The LN parenchyma becomes increasingly segregated as the B cell and T cell zones are established.^7^ While the cortex situated underneath the subcapsular sinus of the LN contains the B cell follicles, T cells and dendritic cells (DCs) interact in the paracortex beneath.^8^ Aside blood endothelial cells (BEC) and LECs, a third major SC population, the reticular fibroblastic stromal cells (FRCs) populate the adult LN. The FRC pool consist of several heterogeneous subsets, including follicular dendritic cells (FDCs), which infrastructurally organize B cell follicles, marginal reticular cells (MRC) below the subcapsular sinus, T cell zone reticular cells (TRC) and medullary FRCs located in the paracortex and the medulla, respectively.^4^ These FRCs are thought to develop from mLTos transitioning through a myofibroblastic precursor stage.^9^ Recent scRNA-seq profiling of LNSCs revealed that the podoplanin (*Pdpn*, gp38)-expressing SC compartment encompasses a distinct and heterogeneous population of non-endothelial CD34^+^ SCs, which are located at the LN capsule or the adventitia of large vessels.^10–12^ While several lineage-tracing models have delineated the prenatal origin of non-endothelial LNSCs,^13–15^ the postnatal differentiation process of FRCs and CD34^+^ SCs, along with their immuno-modulatory functions, presumably tightly regulated by epigenomic modifications and gene regulatory networks, is less well defined.

Although, the initial priming of the adaptive immune response heavily relies on antigen-presenting DCs, the intrinsic microenvironment of the respective tissue-draining LN and its SC compartment, greatly influences this process. Particularly, the heterogeneous population of FRCs has been shown to shape the adaptive immune responses by providing survival molecules such as IL-7 and BAFF,^16,17^ or readily upregulating iNOS upon IFNγ signaling, thereby limiting the expansion of pro-inflammatory T cells and globally suppressing aberrant priming of adaptive immune cell differentiation.^18,19^ Distinct localization of FRC subsets, such as Cxcl9-producing FRCs, can mitigate effective migration of Cxcr3^+^ cells, including memory CD8^+^ T cells, during the course of antiviral immune responses.^12,20^ In addition to these immuno-modulatory functions inherent to any LN, the local SC compartment also tissue-specifically shapes the migratory and effector T cell properties.^11,21,22^ Despite the detailed understanding of prenatal LN development, little is known about which transcriptional regulators impinge postnatally on the presumably common pool of mLTos to give rise to the heterogeneous population of non-endothelial SCs^23^ and their immune-modulatory potential.^11,22,24^ Although multiple mechanisms through which mesenchymal SCs modulate the LN microenvironment have been identified, the underlying transcriptional regulators and epigenomic framework defining distinct functional SC responses from birth and throughout adulthood are only incompletely understood.^25,26^

Here, we have mapped the location-specific epigenomic landscape of non-endothelial SCs encompassing both FRCs and CD34^+^ SCs. We identified location-specific, epigenetically modified genomic loci, and based on motif enrichment analysis together with gene expression profiling, delineated steady-state expression signatures poising peripheral skin-draining LN (pLN) SCs for pro-inflammatory responses. Furthermore, by obtaining the first developmental scRNA-seq profiling dataset following mesenteric lymph node (mLN) SC development from birth throughout adult life, we also observed the postnatal segregation of mesenchymal progenitors and confirmed their distinct localization within the developing mLNs. We further outlined the developmental trajectory of the mLNs’ SC compartment and mapped subset-specifically expressed transcription factors (TF) to distinct expanding SC populations, which shape the epigenomic landscapes across CD34^+^ SCs and FRCs. Finally, using lentiviral overexpression, we uncovered *Irf3* as a key transcriptional regulator driving mesenchymal stem cell differentiation towards Cxcl9^+^FRCs.

Our data constitute a comprehensive transcriptional map of SC development in mLNs. More precisely, we characterize the transcriptional and epigenomic landscape of non-endothelial LNSCs, providing a valuable resource to delineate transcriptional regulators that govern the developing LN and thereby permanently shape adaptive tissue-specific immune responses.

## Results

### Location defines the epigenomic landscape of non-endothelial LNSCs

Previous studies underlined the immuno-modulatory functions of LNSCs.^27^ We thus aimed to delineate epigenomic modifications that underlie the immuno-modulatory function of LNSCs. To this end, we performed whole-genome bisulfite sequencing (WGBS) and assay for transposase accessible chromatin sequencing (ATAC-seq) to identify CpG methylation and genomic accessibility, respectively, of CD31^−^Pdpn^+^ non-endothelial SCs isolated by fluorescence-activated cell sorting (FACS) from mLNs and pLNs originating from adult mice housed under specific pathogen-free (SPF) or germ-free (GF) conditions. We were able to determine the methylation status of 88.2 % of the 2.19*10^7^ CpGs and, using *Bsmooth*,^28^ identifying 1532 non-overlapping differentially methylated regions (DMRs) across all pair-wise comparisons. Importantly the vast majority of the DMRs were location-dependent (**Fig. 1A**), whereas only 16 and 17, were commensal-dependent for pLN and mLN, respectively (**Supplementary Table 1**). Over 390 DMRs, located in the proximity of the transcription start site (TSS), were annotated to 286 genes, including microenvironmental mediators (e.g. *Cxcl13, Gdf6, Sfrp5*), immuno-modulatory enzymes (e.g. *Aldh1a2, Ptgis*),^29–32^ and TFs (e.g. *Isl1, Hoxd1, Meis1, Meis2, Nkx2-3, Tcf4, Foxn2*), potentially impinging on transcriptional regulation (**Fig. 1B-C**).^33^

**Figure 1.**
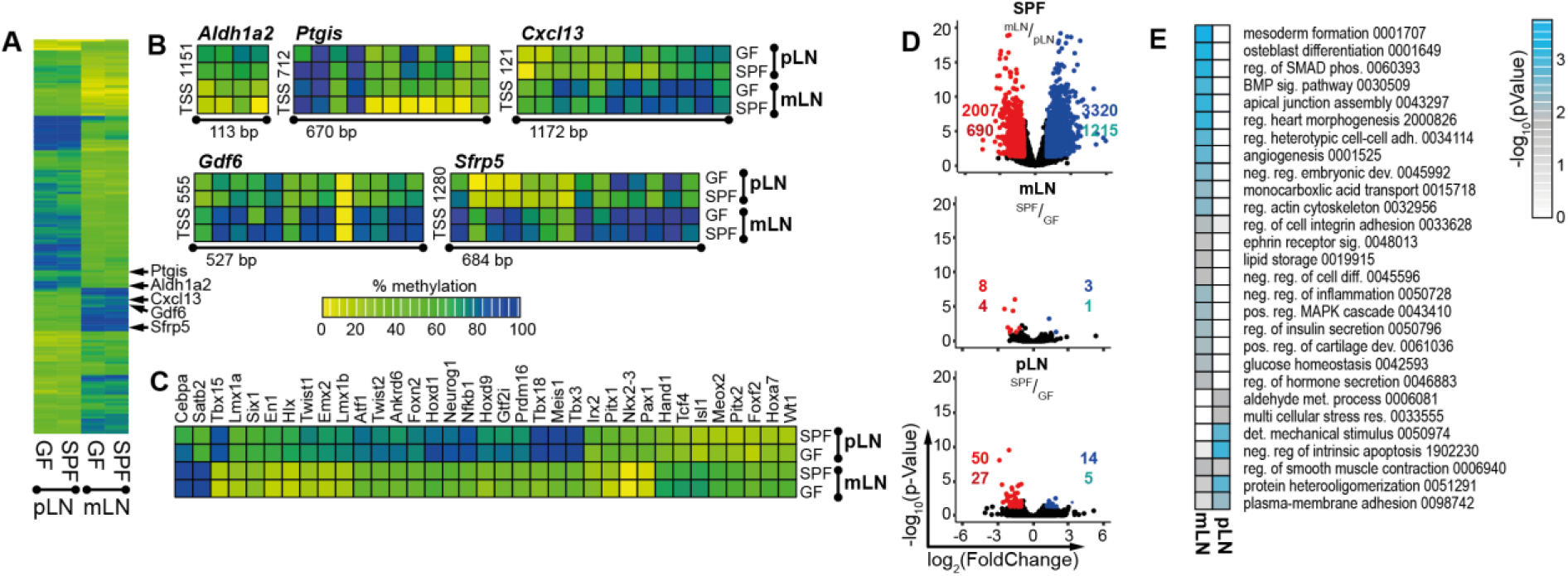
LN location defines the epigenomic landscape of non-endothelial SCs. CD45^−^CD24^−^CD31^−^Pdpn^+^ SCs were isolated from mLNs and pLNs of GF or SPF mice, and WGBS (A-C) or ATAC-seq (D-E) analyses were performed. **(A-C)** DMRs were identified in colonization- (SPF vs. GF) and location-dependent (mLN vs. pLN) pairwise comparisons. Scale bar depicts the extent of methylation with 100 % being fully methylated and 0 % being non-methylated. **(A)** The heatmap represents the mean methylation of significant DMRs within the promotor region of 284 genes. **(B)** Heatmaps represent CpG methylation of exemplary DMRs. The distance from the TSS is indicated and the size of genomic loci denoted in base-pairs (bp). **(C)** The heatmap represents the mean methylation of significant DMRs within the promotor region of TFs. **(D-E)** DARs were identified in colonization- (SPF vs. GF) and location-dependent (mLN vs. pLN) pairwise comparisons. **(D)** Volcano plots of mean ATAC-seq FPKM comparing indicated samples. The number of DARs (top) and genes (bottom) is indicated per comparison. **(E)** GO analysis of biological processes of genes with location-dependent DARs. The numbers denote GO identifiers. ATAC-seq, assay for transposase accessible chromatin sequencing; DAR, differentially accessible region; det, detection; DMR, differentially methylated region; GF, germ-free; GO, gene ontology; met, metabolism; neg, negative; pos, positive; reg, regulation; SC, stromal cell; sig, signalling; SPF, specific pathogen-free; TF, transcription factor; TSS, transcription start site; WGBS, whole-genome bisulfite sequencing.

We then proceeded to obtain a global overview of chromatin accessibility. Peaks were called per sample and merged to obtain common open genomic regions across all conditions comprising a total of 42,434 peaks (**Supplementary Table 2**) As expected, we identified a substantial number of 5327 differentially accessible regions (DARs) between mLN-SPF and pLN-SPF, whereas absence or presence of microbiota only marginally influenced the accessibility profile of non-endothelial SCs for both mLN and pLN, corroborated by the low number of detected DARs (11 and 64, respectively; **Fig. 1D**). Gene ontology (GO) analysis of the location-dependent DARs revealed that mLN SCs are responsive to BMP signalling and negatively regulate inflammatory processes (**Fig. 1E**). Surprisingly, pLN SCs did not show increased chromatin accessibility for pro-inflammatory mediators (**Fig. 1E**), which is in contrast to the previously made observations at the transcriptional level.^11,34^

Altogether, these data implicate that the tissue-specific location of LNs strongly influence the epigenomic landscape of non-endothelial SCs, thereby potentially contributing to TF-controlled transcriptional programs.

### Differential accessibility delineates location-dependent regulators of persistent SC function

The observed discrepancy between gene-associated DARs and previously published gene expression signatures particularly for non-endothelial pLN SCs^11,34^ prompted us to further dissect transcriptional and epigenomic co-regulation. In order to compare both transcriptome and chromatin accessibility, we obtained RNA-seq data of CD31^−^Pdpn^+^ non-endothelial SCs isolated using FACS from mLNs and pLNs originating from adult mice housed under SPF conditions. We then correlated location-dependent differential expression with the respective chromatin accessibility within the promotor region on a per-gene basis, which allowed us to divide genes into three distinct groups (**Fig. 2A**). The first group includes genes whose transcriptional activity correlates with chromatin accessibility, with 207 and 183 for mLN SCs and pLN SCs, respectively. These genes comprise 24 % of the differentially expressed genes (DEGs) and include genes such as *Nkx2-3*, *Ccl20* and *Cxcl9*, known for their location-dependent differential expression.^11,34^ The second group contains 43 % of the regulated genes that are differentially accessible, but not differentially expressed, potentially composed of elements that respond to cellular activation. Interestingly, 296 genes for mLN SCs and 255 genes for pLN SCs are differentially expressed, but have no associated DARs, indicating that these genes are regulated by TFs that exploit already accessible chromatin and that we define here as active TF regulated (**Fig. 2A**). We performed GO analysis for biological processes on the inducible gene modules or active TF regulated genes for mLN SCs and pLN SCs, and identified particularly for the genes associated with active TFs for pLN SCs a substantial enrichment of GO terms associated with elevated immune responses including *neutrophil chemotaxis* and *response to cytokine* (**Fig. 2B-C**). Interestingly, within mLN SCs, GO terms indicative of immune suppression were enriched, including the *negative regulation of Nfkb signalling* (**Fig. 2B-C**).

**Figure 2.**
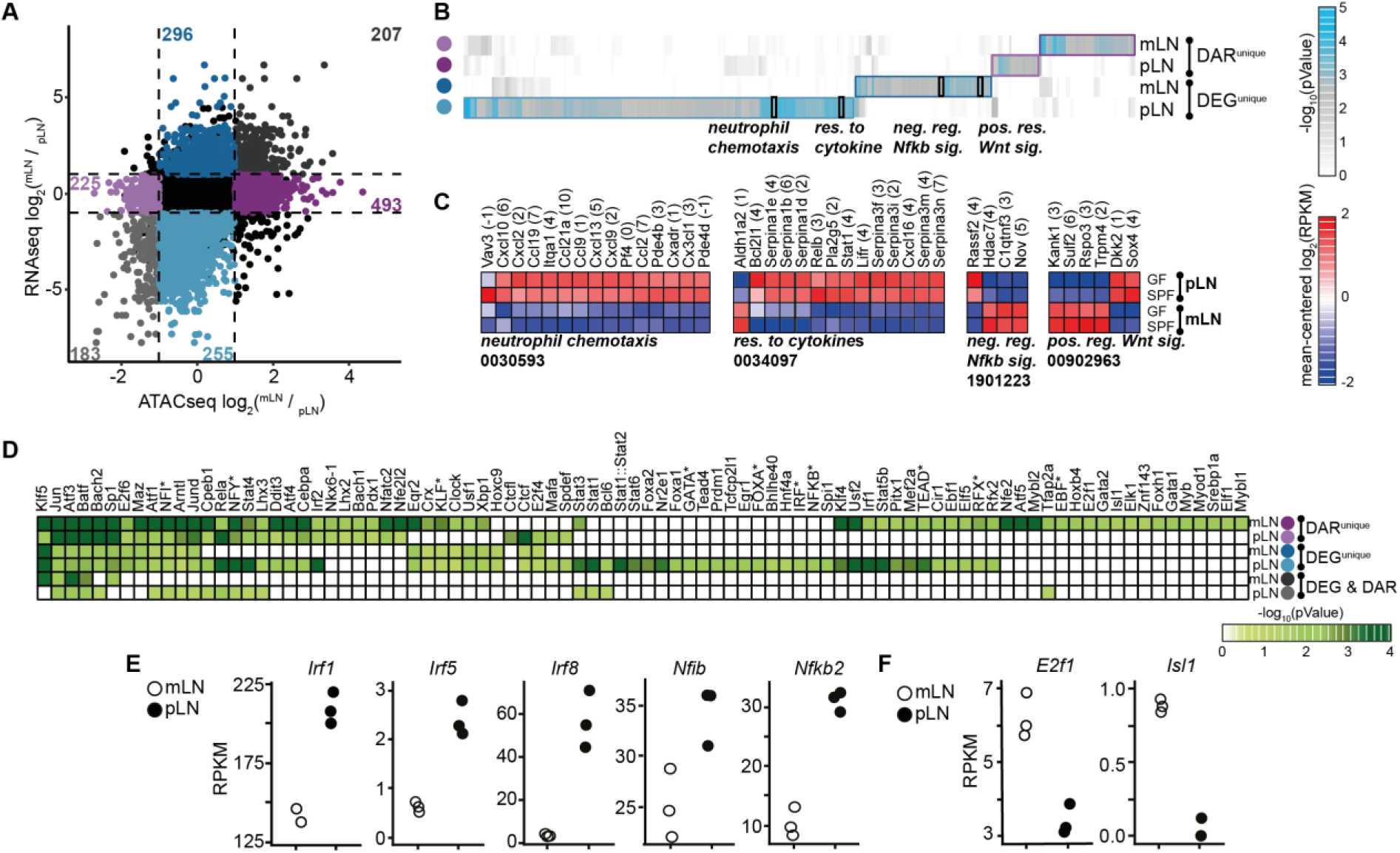
Accessible and actively expressed TFs promote pro-inflammatory gene expression profile of pLN non-endothelial SCs. CD45^−^CD24^−^CD31^−^Pdpn^+^ SCs were isolated from mLNs and pLNs of SPF mice. Subsequently, RNA-seq (A-F) or ATAC-seq analyses (A-B, D) were performed. DEGs and DARs were identified in mLN vs. pLN pairwise comparisons. **(A)** Colored numbers in the scatterplot represent the number of genes with DAR and/or differential expression. Only genes with accessible loci within the promotor region were included in the analysis. On the x-axis, log_2_(FC) of accessibility per DAR is plotted and on the y-axis, the log_2_(FC) of gene expression for the comparison of mLN vs. pLN. **(B)** GO analysis of biological processes of DEGs and/or genes with at least one DAR. **(C)** Heatmaps represent the expression of all DEGs within the GO groups highlighted in (b). The numbers denote GO identifiers. **(D)** The enrichment of known TFBS motifs for each of the quadrants in was utilized to identify TFBS over-represented in the accessible chromatin. The heatmap represents the p-value for enriched TFBS and its corresponding putative TF(s). **(E)** The scatterplots depict differentially expressed TFs that were uniquely identified from the accessible loci of the 255 DEGs for pLN. **(F)** The scatterplots depict differentially expressed TFs that were uniquely identified from the accessible loci of the 493 DEGs for mLN. ATAC-seq, assay for transposase accessible chromatin sequencing; DAR, differentially accessible region; DEG, differentially expressed gene; det, detection; FC, foldchange; SC, stromal cell; GO, gene ontology; neg, negative; pos, positive; reg, regulation; res, response; sig, signalling; TFBS, transcription factor binding site; TF, transcription factor.

Next, we aimed to identify the TFs responsible to alter expression without modifying chromatin accessibility under steady-state conditions. To this end, we identified over-represented TF binding sites (TFBS) and their putative TFs, for the inducible gene modules and active TF regulated genes within accessible genomic loci of the respective genes. As expected, a substantial number of TFs, namely *Klf5, Jun, Atf3, Batf, Bach2, Sp1, Atf1, Nf1* and *Jund* were enriched in at least five out of six gene loci sets indicating their general involvement in shaping the function of non-endothelial SCs (**Fig. 2D, Supplementary Fig. 1**). Surprisingly, only the inducible gene module for mLN SCs and the active TF regulated module for pLN SCs showed unique over-representation of TFBS and putative TFs (**Fig. 2D**). Therefore, we assessed which of the putative TFs identified from the genomic loci within pLN SCs are differentially expressed, and determined *Irf1, Irf5, Irf8, Nfib* and *Nfkb2* to potentially contribute to the increased transcription of pro-inflammatory genes in pLN SCs under steady-state conditions (**Fig. 2E**), whereas *E2f1* and *Isl1* were solely identified from inducible gene modules of mLN SCs (**Fig. 2F**).

Together with a large number of shared TFs, the observed TFBS enrichment patterns suggest that non-endothelial SCs contain a location-dependent epigenomic landscape, allowing distinct TF regulation in pLN and mLN. Our results moreover uncovered a defined set of TFs, including *Irf1, Irf5* and *Irf8*, that drive increased expression of pro-inflammatory cytokines in pLN non-endothelial SCs independent of chromatin accessibility and pinpoint uniquely active TFs in pLN and mLN.

### Developmental age constrains cellular and functional composition of the mLN SCs

We hypothesized that the observed differences across the epigenomic landscapes of non-endothelial SCs derive from epigenomic modifications that were obtained during early postnatal development. It is widely accepted that substantial epigenomic modifications occur during the transition of progenitors to fully differentiated cells.^35,36^ As LNs undergo rapid postnatal expansion, we first aimed to dissect changes in SC composition along mLN development.

To this end, we resected mLNs from mice at early postnatal age (day 0/1, D0), the early (D10) and late (D24) juvenile stages, adult age (D56), as well as old age (D300), and obtained FACS-purified CD45^−^CD24^−^ cells. We performed single-cell (sc)RNA sequencing and gathered transcriptomes of 15,925 cells with comparable sequencing depth (55,000-98,000 mean reads per cell) across the different time points. Initially, we used transcriptional signatures for mLN SCs^11^ and the canonical marker for endothelial cells *Pecam1* (**Supplementary Fig. 2A**) to remove endothelial cells and perivascular cells (PvC) from the analysis, after alignment of samples using diagonal canonical correlation analysis (Methods). Additionally, we observed and excluded cells that were disproportionately represented in D0/1 mLNs, expressing significantly higher levels of the growth factors *Igf1* and *Igf2* as well as the extracellular matrix protein *Mfap4*, indicative of adjacent tissue fibroblasts (**Supplementary Fig. 2B-C, Supplementary Table 3**). To obtain an unbiased picture, we re-embedded 5,658 non-endothelial SCs across all five developmental stages. Twelve transcriptional clusters harboring unique functional properties were identified based on DEGs, GO analysis and previously published signatures (**Fig. 3A, Supplementary Fig. 2D-E, Supplementary Table 4**).^11,12^ These clusters were broadly separated into CD34^+^ SC, including CD34^+(CD248+)^, CD34^+(Ackr3+)^, CD34^+(Aldh1a2+)^, metabolically active (mFRC), Il6^+(Cxcl1+)^ FRC, Cxcl9^+^ FRC, Inmt^+^ FRC, Inmt^+(Cxcl12+)^ FRC and Ccl19^+(Il7+)^ FRC, mesothelial-like (Meso), LTo-like and progenitor-like (Prog) subsets (**Fig. 3A**). All identified clusters were variably represented along postnatal mLN development (**Fig. 3B-C**), with particularly LTo-like and Prog subsets being over-represented at D0 and D10. As expected, LTo-like SCs highly expressed *Cxcl13*, *Tnfsf11* and *Madcam1* (**Fig. 3D-E**).^2,3,9^ Interestingly, we observed an additional highly proliferating subset, here termed postnatal progenitors (Prog; **Fig. 3C-E**). The progenitor subset could further be subdivided into two distinct populations (**Fig. 3F**). Prog^Cxcl13+^ cells expressed significantly higher levels of canonical markers for LTo-like cells including *Madcam1* and *Ccl19*, indicative of a close relation to previously described LTos and their propensity to function as progenitors during LN development.^37^ Surprisingly, we identified an additional subset, here termed Prog^CD34+^ which showed higher expression for several collagens (*Col3a1, Col6a2* and *Col1a2*), as well as genes involved in regulating Bmp-signalling including *Gpc3* and *Crip* (**Fig. 3F**).^38^

**Figure 3.**
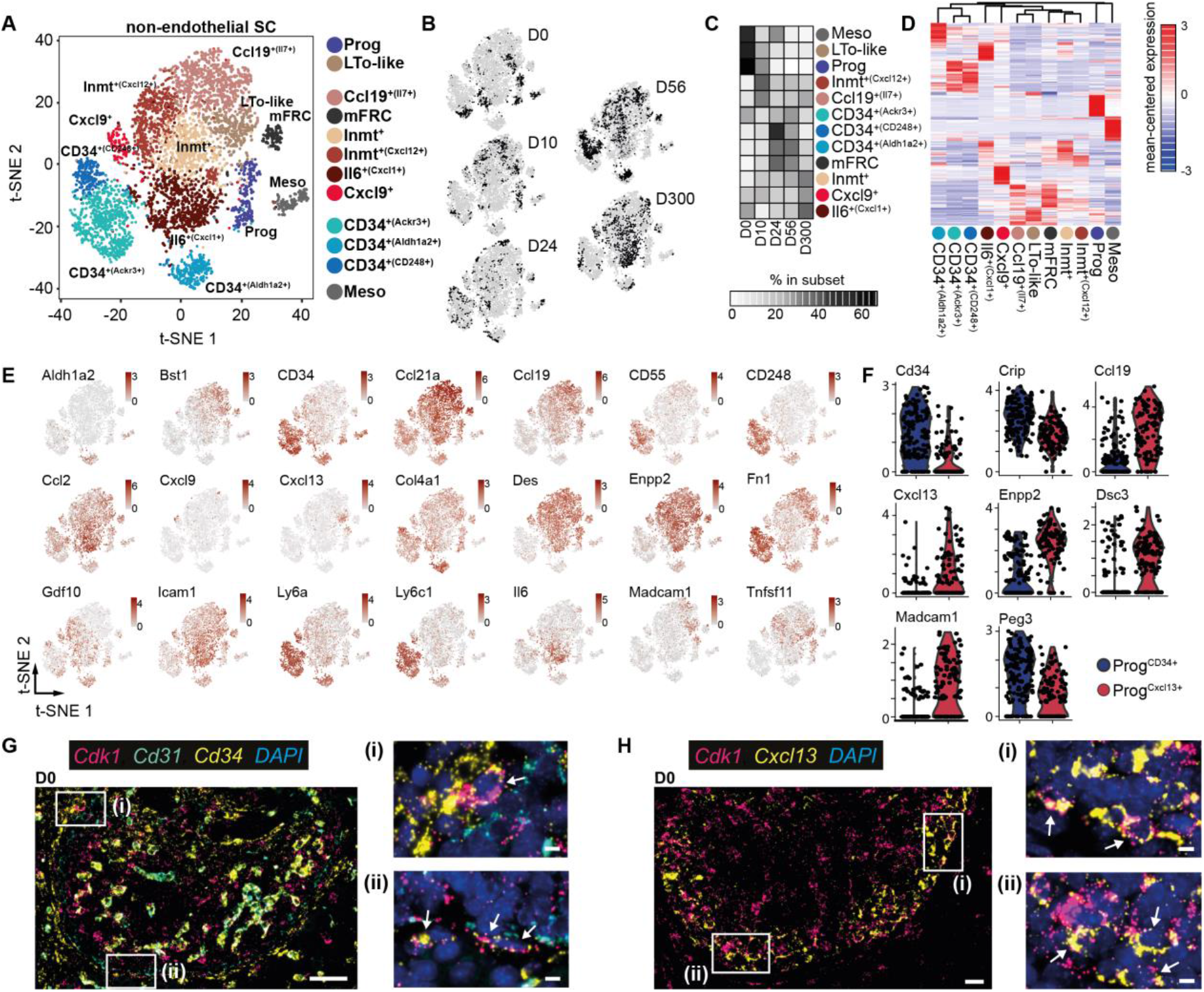
Postnatal ontogeny of non-endothelial mLN SCs. CD45^−^CD24^−^ cells were isolated from mLNs of day 0, 10, 24, 56 and 300 old SPF-housed mice and subjected to scRNA-seq. Non-endothelial SCs were identified as non-LECs, non-BECs and non-PvCs. **(A)** t-SNE plot of merged SCs across ages showing cluster segregation. **(B)** t-SNE plot of SCs from each age. **(C)** The heatmap represents the percentage of cells in each subset across time points normalized to cell number. **(D)** Hierarchical clustering of subsets based on the mean expression of the top 40 DEGs per subset. **(E)** t-SNE plot colored for expression of segregating genes. **(F)** Violin plot of DEGs for subsets identified among putative postnatal progenitors. **(G**-**H)** Sections (3 μm) of D0 neonatal mLN were stained with the indicated RNAscope probes and imaged by fluorescence microscopy. Nuclei were counter-stained with DAPI (blue). Images represent an overview of mLN anlagen with “white squares” indicating regions of interest. ***(i-ii)*** Images represent zoom-ins of respective regions of interest. Arrows indicate marker co-expression per cell. Representative tissue sections n = 2-3. **(G)** Capsular positioning of *Cd34^+^Cdk1^+^* cells. Overview (scale bar = 50 μm) and zoom-in (scale bar = 5 μm) for RNA probes specific to *Cdk1, Cd31* and *Cd34*. **(H)** Cortex-localized *Cxcl13^+^Cdk1^+^* cells. Overview (scale bar = 20 μm) and zoom-in (scale bars = 5 μm) for RNA-probes specific to *Cdk1* and *Cxcl13*. BEC, blood endothelial cell; LEC, lymphatic endothelial cell; LTo-like, lymphoid tissue organizer like cell; Meso, mesothelial-like cells; mFRC, metabolically active FRC; Prog, postnatal progenitor; PvC, perivascular cell; SC, stromal cell.

We then aimed to identify the localization of the respective putative progenitor populations within the developing LNs by utilizing RNAscope on neonatal mLN tissue slices. Initially, we confirmed that i) the RNA integrity of D0 mLN remained intact after resection by staining for a mix of ubiquitously expressed genes (*Polr2a, Ubc, Ppib*) as a positive control (**Supplementary Fig. 3A**) and ii) the majority of *Ccl19* and *Cxcl13* expressing cells were spatially separated from *Cd34^+^* cells at this stage of postnatal mLN development (**Supplementary Fig. 3B-C**). Further, we utilized cyclin-dependent kinase 1 (*Cdk1*), a key kinase at the transition to the S-phase, which is highly expressed in both of the two putative progenitor populations, as a proxy to pinpoint progenitor cells. We readily identified capsular *Cd34* and *Cdk1* expressing non-endothelial *Cd31*^−^ cells (**Fig. 3G**). Importantly, *Cd34*^+^*Cdk1*^+^ cells were neither present within the cortical nor paracortical areas, but could be identified within adjacent tissue (**Supplementary Fig. 3D**). As expected, we furtheron detected *Cdk1^+^Cxcl13^+^* cells,^3,4^ predominantly in the developing cortex (**Fig. 3H**).

Together, these data delineate the postnatal expansion of the heterogeneous non-endothelial SC compartment and identify two putative postnatal progenitor cell populations that are distinctly positioned within the developing LN.

### Distinct progenitors establish the non-endothelial SC compartment rapidly after birth

Previous studies showed that postnatal expansion of non-endothelial SCs within the LN relies on the progenitor potential of LTo cells, differentiating into various SC subsets.^14,39,40^ Additionally, progenitors with LNSC potential have been shown to reside in the CD34^+^ perivascular niche of multiple adult organs.10 Therefore, we aimed to elucidate whether LTo-like, putative Prog^Cxcl13+^ and Prog^CD34+^ progenitors observed at early postnatal mLN development have the molecular potential to give rise to CD34^+^ SC and FRCs in the expanding neonatal mLN.

We initially constructed a trajectory using all non-endothelial SCs, and observed two distinct sets of branches evolving along physiological development (**Supplementary Fig. 4A**), underscored by expression of key marker genes identifying progenitors at distinct, presumably early stages of development (**Supplementary Fig 4B**).^27,37,41^ We therefore re-embedded cells situated along the CD34^+^ SC or FRC trajectory (**Supplementary Fig. 4C**) and employed pseudo-time mapping using Monocle2.^42^ As expected, early developmental time points corresponded to pseudotemporal branches at the starting point of the trajectory (**Fig. 4A-B**). Importantly, already at D24, CD34^+^ SCs were distributed along the complete pseudotemproal space, whereas only minor changes in the distribution of cells across pseudotime were observed beyond D56, indicating that main non-endothelial SC differentiation is established shortly after weaning (**Fig. 4A-B**). While *Ccl19* and *Ptgis* were constitutively expressed over the developmental trajectory, subset-defining genes including *Aldh1a2* and *Il6* were predominantly expressed at distinct branches of the CD34^+^ SC or FRC trajectories, respectively (**Fig. 4B-C**), indicative of developmental segregation for each subset of CD34^+^ SCs and FRCs, culminating in distinct functional subsets (**Fig. 4D**). Importantly, *Cxcl13* was expressed highest early during development in FRCs (**Fig. 4C**). As expected, pseudotime starting points of both the CD34^+^ SC and FRC trajectories comprised predominantly Prog^CD34+^ or Prog^Cxcl13+^ cells (**Fig. 4E**). Within the FRC trajectory, Prog^Cxcl13+^ cells pseudotemporally aligned with the LTo-like ones, indicating that these cells are of similar origin, giving rise to two major branches of differentiation that correspond to different subsets of FRCs, namely Ccl19^+(Il7+)^ and Il6^+(Cxcl1+)^ (**Fig. 4E-F**). Distinct differentiation paths were underscored by an increased expression of *Tnfsf13b, Il33, Ccl19* and *Tgfbi* for Ccl19^+(Il7+)^, and *Cxcl1* and *Il6* for Il6^+(Cxcl1+)^ subsets along pseudotemporal development (**Fig. 4C, Supplementary Fig. 4D**).

**Figure 4.**
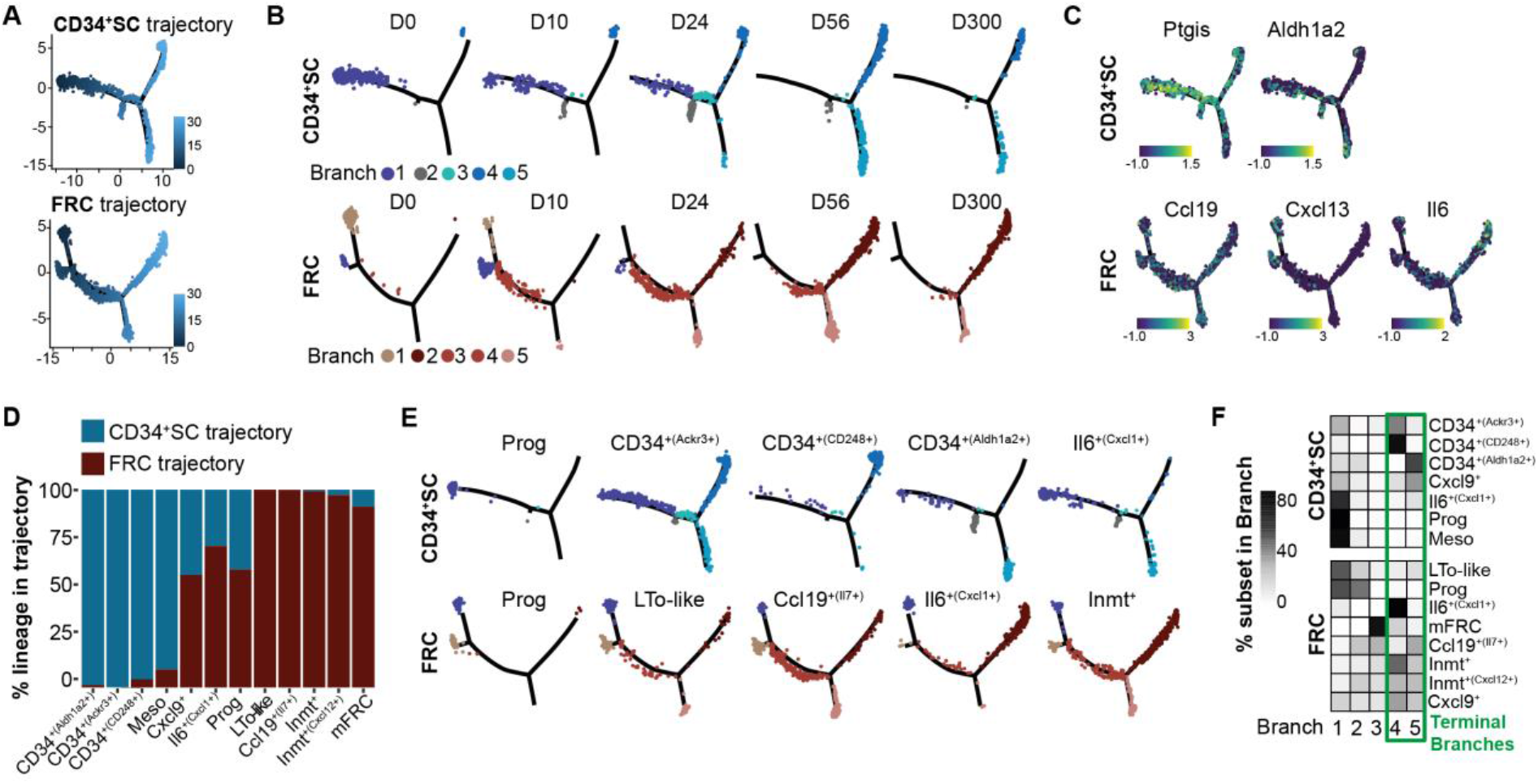
Distinct postnatal progenitors give rise to FRCs and CD34^+^ SCs. CD45^−^CD24^−^ cells were isolated from mLNs of day 0, 10, 24, 56 and 300 old SPF-housed mice and subjected to scRNA-seq. Non-endothelial SCs were identified as non-LECs, non-BECs and non-PvCs. **(A)** Pseudotime ordering of FRCs and CD34^+^ SCs. **(B)** Pseudotime trajectories superimposed with cells per time point. **(C)** Gene expression on pseudotime trajectories. **(D)** Bar graph depicts the proportion of cells from the indicated subset within the FRC or CD34^+^ SC trajectory. **(E)** Cells per cell subset across branches superimposed on pseudotime trajectories. **(F)** The heatmap represents the percentage of cell subsets across branches normalized to cell number. BEC, blood endothelial cell; FRC, reticular fibroblastic stromal cell; LEC, lymphatic endothelial cell; mLN, mesenteric lymph node; PvC, perivascular cell; SC, stromal cell; SPF, specific pathogen-free.

In line with the FRC trajectory, the CD34^+^ SC development initiated from Prog^CD34+^ cells, branching into two major axes consisting predominantly of either CD34^+(CD248+)^ or CD34^+(Aldh1a2+)^ (**Fig. 4E-F**). As expected, a small proportion of CD34^+^ SCs,^2,10^ was already in place at D0, rapidly expanding at D10 and D24 (**Fig 4B**). As anticipated, based on the heterogenous composition of the mLN’s non-endothelial SC compartment, the main terminal branches showed distinct expression profiles. While the CD34^+(CD248+)^ branch expressed higher levels of *Ptgs1* and *Ptgis* enabling prostacyclin synthesis (**Fig. 4C, Supplementary Fig 4D**), the CD34^+(Aldh1a2+)^ branch expressed higher levels of *Col15a1* and *Vtn* together with *Fgf7* and *Gdf10* potentially generating a specific cell adhesion environment (**Supplementary Fig. 4D**). Surprisingly, cells of the Cxcl9^+^ subset were distributed similarly between FRCs and CD34^+^ SCs and were not over-represented at a specific terminal branch, neither in the FRC nor the CD34^+^ SC trajectory (**Fig. 4D**). The dispersion of the Cxcl9^+^ subset across the two main subsets and the trajectory is underscored by the unique expression of *Ly6a*, also a core feature of CD34^+^ SCs, but the lack of consistent *Cd34* expression (**Fig. 3E**).

In sum, the pseudotemporal dissection of the non-endothelial SC compartment indicates that postnatal expansion of the main populations of mLN SCs is driven by two proliferating progenitor populations, both of which are already postnatally set on a defined FRC or CD34^+^ SC differentiation trajectory.

### TFs shape the subset-specific epigenomic landscape early during ontogeny

TFs guide epigenomic modifications to respective target loci, particularly during differentiation processes.^43–45^ We hypothesized that most TFs that drive the differentiation of the two main non-endothelial SC subsets from progenitors to fully functional CD34^+^ SCs or FRCs, as inferred from footprinting analyses of the accessible chromatin, should also be dynamically expressed along mLN development. To assess this, we explored TF binding motif enrichment in the epigenomic landscape of mLN SCs and investigated which TFs are expressed at key branching points of the FRC and CD34^+^ SC differentiation trajectories.

To this end, we combined the over-represented TFBS and their putative TFs, within accessible genomic loci (**Fig. 2D**) with the trajectoral expression of all TFs detected in the developmental scRNA-seq analysis of CD34^+^ SCs and FRCs (**Fig. 4**). In total, 30% of the TFs, identified from the accessible chromatin, were differentially expressed over the developmental trajectory, of which 7 and 13 TFs were uniquely differentially expressed at the key branching point for CD34^+^ SCs and FRCs, respectively (**Fig. 5A-C**). Particularly, the commonly over-represented TFs (**Fig. 2D**) are expressed at the key branching point of both CD34^+^ SC and FRCs (**Fig. 5A-B**), including *Atf3, Egr1, Egr2, Irf1, Jun, Jund* and *Klf9*, indicating that these TFs shape the general course of chromatin accessibility of differentiating SCs (**Fig. 5A-C, Supplementary Fig. 4E**). Importantly, 40% of the TFs that were differentially expressed and motif enriched TFs between pLN SCs and mLN SCs are shared between both CD34^+^ SCs and FRCs, indicating their general requirement in shaping the epigenomic landscape and function of these two non-endothelial main SC subsets (**Fig. 5C**).

**Figure 5.**
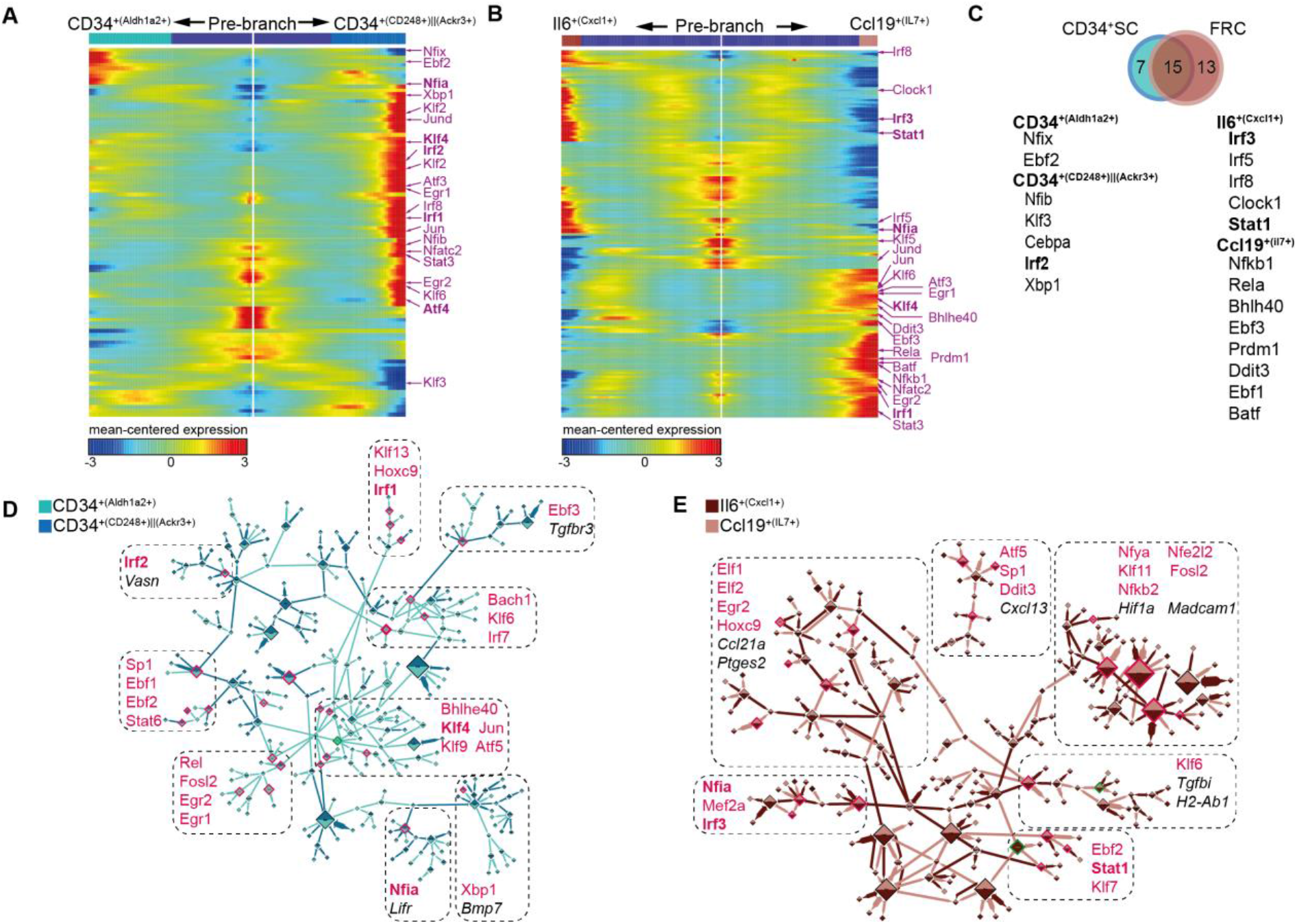
TFBS accessibility aligns with dynamically regulated TFs to shape differentiation and function of key mLN SC subsets. **(A-B)** Heatmap showing expression of TFs over pseudotime involved in the differentiation of **(A)** CD34^+^ SCs and **(B)** FRCs. TFs highlighted in “pink” are derived from TFBS enrichment analyses of mLN SC DARs. **(C)** The Venn diagram depicts the overlap of accessibility-derived TFs. TFs upregulated per subset/branch are denoted in “black”. **(D-E)** Gene regulatory networks based on genes that are dynamically co-expressed along the developmental trajectory, ordering cells within each “branch” per age (see Figure 4). The size of the node is proportional to the node outdegree. TFs inferred from their respective TFBS in accessible chromatin regions (see Figure 2) annotated to the respective node cohort in „pink‟ | DARs and “black” | SC-associated genes. The color of the edges signifies the terminal branch from which the edge was derived. **(D)** depicts CD34^+^ SCs. **(E)** depicts FRCs. DAR, differentially accessible region; det, detection; DMR, differentially methylated region; FRC, reticular fibroblastic stromal cell; mLN, mesenteric lymph node; SC, stromal cell; TF, transcription; TFBS, transcription factor binding sites.

We next utilized *dynGENIE3*^46^ to identify key TFs that potentially impinge upon the differentiation of the cellular components of the expanding mLN, and constructed a gene regulatory network of dynamically regulated genes encompassing TFs for each of the two terminal branches for both FRCs and CD34^+^ SCs. In line with the branch-point-based analysis, we observed that 23% of the dynamically regulated TFs were co-regulated consistently across FRCs and CD34^+^ SCs, including *Nfia*, *Egr2* and *Ebf2* (**Fig. 5D-E**). Despite the substantial proportion of common TF denominators, particularly for the IRF and KLF/SP TF family, a noticeable number of TFs were uniquely regulated along the trajectories. While *Irf1* and *Klf4*/*9*/*13* emerged for CD34^+^ SCs, *Irf3* and *Klf6*/*7*/*11* were solely identified for FRCs (**Fig. 5C-E**), suggesting dissimilar modulatory paths during the development of these two distinct non-endothelial SCs.

In summary, by combining branch-point-based differential expression together with identification of dynamically co-regulated gene networks, we were able to identify TFs that are potentially decisive in postnatally shaping the differentiation of CD34^+^ SCs or FRCs.

### Members of the IRF transcription factor family contribute to the differentiation of CD34^+^ SCs and FRCs

By combining orthogonal analyses of TFBS enrichment and gene expression along mLN development, we were able to identify a concise assembly of TFs that are putatively involved in the differentiation of non-endothelial SC subsets. We therefore hypothesized that these TFs should be able to drive the differentiation from multipotent progenitor cells towards a cell-type or state-specification resembling *ex vivo* profiled SC subsets.

To this end, we utilized a well-established TF-overexpression model based on lentiviral integration, puromycin selection and timed doxycycline-driven overexpression^47^ in a murine C3H10T1/2 mesenchymal stem cell like (MSC) line.^48^ We focused on TFs that were either identified for both CD34^+^ SCs and FRCs (e.g. *Atf4, Nfia, Irf1*), only for CD34^+^ SCs (e.g. *Irf2, Cebpa*), or only for FRCs (e.g. *Irf3*, *Stat1*). We also included *Myc*, as LTos have myofibroblastic features,^9^ and *Fos* as it is widely expressed across all non-endothelial SC (**Supplementary Fig. 2E**).^11,12^ We initially assessed cellular morphological changes of differentiated MSCs and observed a broad range of morphologies, including upregulation of lipid droplet formation for *Nfia*^49^ and distinctive fibroblastic features for the majority of overexpressed TFs, including *Irf1, Irf3* and *Myc* (**Fig. 6A**). We then assessed the gene expression profile of induced MSCs by performing bulk 3’RNA-seq (BRB-seq) at the terminal differentiation time point, 12 days post doxycycline-driven overexpression.^50^ To compare the extent of matching expression signatures for each of the overexpressed TFs with the endogenous subset-specific signatures, we calculated the cumulative Z-score for the Top 100 DEGs for each of the endogenously identified SC subsets (**Supplementary Table 4**). As expected, *Myc* overexpression drove an LTo-like expression signature, consistent with a myofibroblastic origin (**Fig. 6B**).^9^ The most striking overlap of the endogenous expression signature was observed for *Irf3* overexpression, supporting the expression signature of DEGs identified for the Cxcl9^+^ FRCs, which was further corroborated on a per cell basis when calculating the cZscore from the *Irf3-*driven overexpression (**Fig. 6C**). Furthermore, *Irf3* overexpression supported the upregulation of Cxcl9^+^ FRC subset-associated genes, including several interferon-induced proteins (e.g. *Ifit1, Ifit3, Ifit3b*) and antiviral response elements (e.g. *Isg15, Oasl2*) above the expression levels detected for any of the overexpressed TFs including *Irf1* and *Irf2* (**Fig. 6D**).

**Figure 6.**
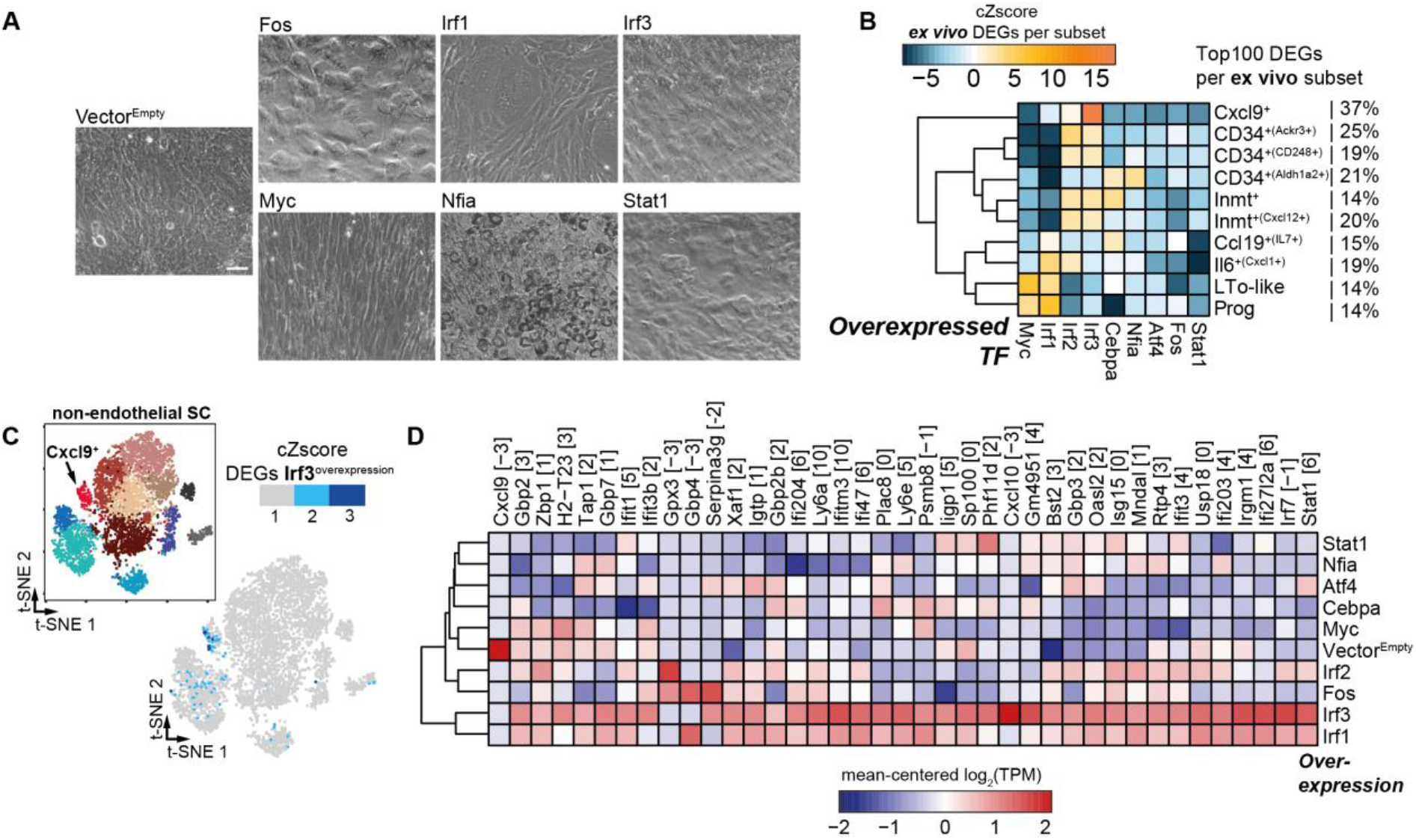
*Irf3* promotes the differentiation of mesenchymal stem cells towards a Cxcl9^+^ FRC molecular phenotype. Murine C3H10T1/2 were lentivirally transduced and puromycin-selected for stable vector integration. TF expression was doxycycline-induced and maintained for 12 days. Subsequently, BRB-seq was performed on the induced cells. **(A)** Representative phase contrast images of morphological phenotypes subsequent to overexpression-driven differentiation (scale bar = 30 μm). **(B)** The heatmap depicts the cumulative Z-score (cZscore) of DEGs, compared to Vector^Empty^, at “TF overexpression” in the C3H10T1/2 cell line among the Top 100 DEGs per LN subset (Supp. Table S4). The percentage indicates the proportion of DEGs that were identified per subset (see Figure 3, Supp. Table S4) and the DEGs identified with “TF overexpression” in C3H10T1/2. (**C)** The cumulative Z-score on a per cell basis of the significant DEGs due to *Irf3* overexpression superimposed on the scRNA-seq developmental map. (**D)** The heatmap depicts all detected DEGs from the Cxcl9^+^ cluster for the tested (overexpressed) TFs. cZscore, cumulative Z-score; DEG, differentially expressed gene; TF, transcription factor; FRC, reticular fibroblastic stromal cell; SC, stromal cell; TPM, transcripts per kilobase of exon per million reads.

Based on the comprehensive epigenomic and gene expression analyses of mLN SC development, we identified TFs that may shape the postnatal immuno-modulatory specialization of LNSCs. By overexpressing a selected set of delineated TFs, we validated *Irf3* to distinctly promote the differentiation from MSCs to a cell phenotype that molecularly resembles Cxcl9^+^ FRCs.

## Discussion

In our study, we set out to map the epigenomic landscape of the non-endothelial SC compartment of both skin- and gut-draining LNs. Together with scRNA-seq profiling along postnatal development, we increased our understanding of four key aspects of the LNs non-endothelial SC compartment: 1) LNSCs are marked by a tissue-specific, microbiota-independent epigenomic landscape; 2) within pLN SCs, defined TFs propel transcriptional upregulation of pro-inflammatory cytokines; 3) Postnatal segregation of mesenchymal progenitors is driving the expansion of the developing LN; and 4) Cxcl9^+^ FRC differentiation is driven predominantly by *Irf3*.

LNSCs have long been recognized as key structural organizers, but are also increasingly perceived as effective modulators of immune responses,^8,23^ even in a location-specific manner.^11,21,22^ However, whether tissue-specific immuno-modulatory properties of the non-endothelial SC compartment are retained in the epigenomic landscape is so far understudied.^25^ Importantly, when we compared both chromatin accessibility and DNA methylation using ATAC-seq and WGBS, respectively, we observed striking differences between skin- and gut-draining LNs, yet we could exclude a key contribution by microbial colonization. Although several studies have shown that the microbiota can impinge on the epigenome of the host’s adaptive immune system,^51^ its influence does not seem to extend to the draining LNs SCs. This finding was surprising, as we had previously demonstrated that mLNs from GF mice lose their high Treg-inducing capacity upon transplantation into a skin-draining site, while the mLNs from SPF-housed mice persistently maintained their immuno-modulatory functions.^11,22^ Hence, the precise epigenomic modifications and the involved SC subsets that enable microbiota-dependent imprinting of the differential Treg-inducing capacity remain to be identified, a feat that should be attainable, considering the increasing sensitivity of methods to profile histone modifications and TF binding,^52,53^ that align with the low cellular yields of *ex vivo* SCs. We would like to point out that the currently applied sorting of CD45^−^CD31^−^gp38^+^ cells, also applied for the epigenomic profiling in the presented study, yields a heterogeneous cell population that could very well mask epigenomic modifications on a per subset basis. Hence, future studies should consider to at least distinguish between the two dominant SC types in the mLN, namely FRCs and CD34^+^ SCs.

Previous studies underlined the tissue-specific immuno-modulatory functions of LNSCs, with the skin-draining pLN SCs tending to reside in a more pro-inflammatory state as compared to mLN SCs.^11,34,54^ We thus aimed to delineate epigenomic modifications that maintain the immuno-modulatory function of LNSCs. By performing motif enrichment analysis together with gene expression profiling, we delineated steady-state expression signatures poising pLN SCs for pro-inflammatory responses and identified TF modules that likely regulate the expression of dormant accessible genes in mLN SCs. The higher expression of pro-inflammatory mediators aligns with previous studies,^11,34^ but the observations here point towards an active TF-based regulation of pro-inflammatory responses in pLN SCs that relies on chromatin features that are equally accessible across gut- and skin-draining LNSCs. The distinctly higher expression of several interferon regulatory factors, including *Irf1, Irf5* and *Irf8* under steady-state in pLN SCs, thus enables the skin-draining LNs to rapidly respond to infectious incursions. Together with the elevated steady-state expression of pro-inflammatory chemotactic mediators, this enforces the propensity of the pLN SC compartment to support pro-inflammatory responses; an epigenetically imprinted feature not maintained within the mLN SCs.

We initially set out to map postnatal SC development to identify TFs that drive the differentiation of distinct subsets and correspond to the epigenomic TF footprint. scRNA-seq profiling along postnatal development identified two distinct subsets of putative proliferating progenitors that harbor the potential to yield either FRCs or CD34^+^ SCs. The surprising observation of distinct postnatal SC subsets with progenitor potential, extends the perception that Cxcl13^+^ LTos are the sole subset that contributes to the expanding non-endothelial SC pool.^8,23,41,55^ Although we cannot formally exclude that CD34^+^ Prog cells originate from Cxcl13^+^ LTos and can not make a detailed statement regarding the precise Cxcl13^+^ LTo and Cxcl13^+^ Prog progenitor-progeny relationship, the inferred differentiation trajectories underscored the distinct gene expression profiles between Cxcl13^+^ Prog and CD34^+^ Prog cells, supporting the notion that these subsets give rise to either FRCs or CD34^+^ SCs. It is conceivable that Cxcl13^+^ Prog cells within mLNs have already established a distinct branch at birth, suggesting that a proliferating subset of commonly identified Cxcl13^+^ LTos is contributing to the postnatal expansion of the LN by transitioning through the Cxcl13^+^ Prog state. In contrast, *Cd34* is expressed in three different locations in the neonatal mLN parenchyma: the capsule, the medulla and the adventitia of major vessels.^10,12^ We noted that large vessels surrounded by an adventitial layer are scarce within the postnatal mLN, thus making it unlikely that the majority of the CD34^+^ cells are of adventitial origin, which is mirrored by the distinct subset-specific gene expression profiles. However, to truly delineate the contribution of different non-endothelial progenitor pools, a spatio-temporal mapping of developing mLNs at pre- and postnatal stages would be required.

Regardless, the postnatal scRNA-seq profiling already enabled us to identify putative progenitor pools and TFs that contribute to the differentiation of distinct subsets of non-endothelial SC in adult mLNs. The identfication of those TFs enables targeted *in vivo* and *in vitro* verification of the determinants that drive the postnatal development of non-endothelial SCs. As none of the canonical markers for distinct subsets of the CD34^+^ SCs or FRCs trajectory are uniquely expressed on a per subset basis, utilization of mouse models involving floxed TFs in conjunction with CD34- or Ccl19-Cre would target larger and less fine-grained subsets of the LN SC compartment. We therefore opted for an *in vitro* TF overexpression model using a MSC line, which enabled us to screen multiple TFs and avoided the requirement to utilize neonatal progenitor cells, which are scarce and can so far not be isolated due to the lack of canonical markers. The majority of the overexpressed TFs did not induce a differentiation towards a phenotypical profile that resembles *ex vivo* SCs, although a substantial number of TFs supported fibroblastic morphology. While the utilized C3H10T1/2 cell line is pre-disposed towards an adipogenic, osteogenic, myogenic or chondrogenic differentiation,^58^ we did observe that overexpression of TFs from the IRF family induced a fibroblastic phenotype, likely due to the embryonic origin and thus multi-potency of the utilized MSCs.^59^ Particularly striking was the observation that *Irf3* drives the differentiation of cells towards a molecular phenotype that resembles Cxcl9^+^ FRCs, while *Irf1/Irf2* support this specific molecular phenotype to a lesser extent. Within the LN, expression of *Cxcl9* by Cxcl9^+^ FRCs is critically required to establish the chemotactic driving forces that enable the initial detection of pathogenic incursions.^20^ The distinct positioning of the proportionally small cellular compartment of Cxcl9^+^ FRCs in close proximity to lymphatic entry points enables rapid detection of tissue-originated pathogenic incursions, and subsequently accumulation of memory CD8^+^ T cells.^12,20^ Importantly, Cxcl9^+^ FRCs are conserved across different LNs including skin- and gut-draining tissues.^11,12^ *Irf3* overexpression established robust upregulation of various interferon response elements, in line with the up-regulation of *Stat1*, but did not result in an elevated expression of *Cxcl9*, a process which could rely either on additional stimulators such as IFNβ or the requirement of combinatorial TF expression.^60,61^ Surprisingly, the Cxcl9^+^ FRCs could neither be annotated to the CD34^+^ SC nor the FRC developmental trajectories and showed varying expression for *Ccl19*, *Ly6c1* and *Bst1*, indicating that Cxcl9^+^ FRCs are still heterogeneous themselves.

We anticipate that spatial- and subset-specific transcriptional and epigenomic profiling applied to non-endothelial SCs will further identify defining features of the immuno-modulatory potential of the heterogeneous non-endothelial SCs of mLNs. Nevertheless, the transcriptomic analysis of the postnatally developing mLN together with the epigenomic profiling across skin- and gut-draining LNs, can function as a guiding resource to dissect LNSC expansion and immuno-modulation; paving the way to identify entry points to tissue-specifically modulate immune responses by leveraging the embedded SC immune memory and unique functional properties.

---

**Supplementary Figure 1.**
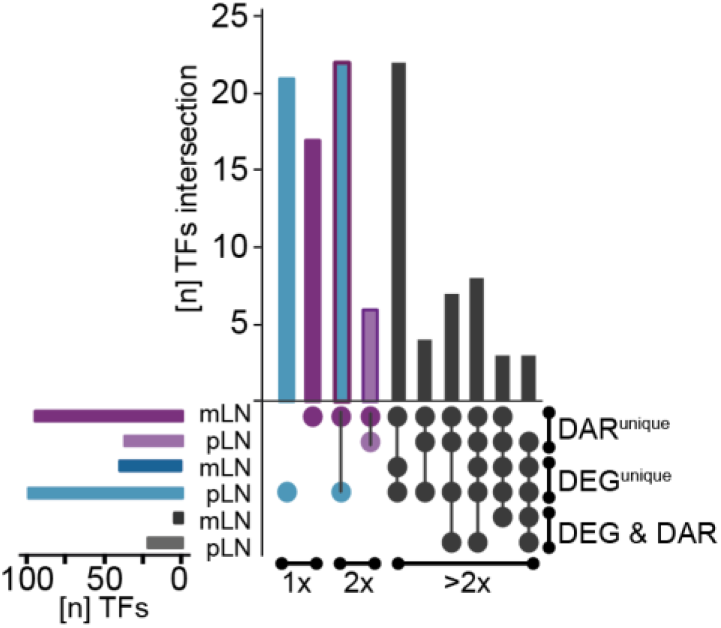
Overlap of TFs identified from the epigenomic landscape and transcribed genes. *Related to Figure 2.* CD45^−^CD24^−^CD31^−^Pdpn^+^ SCs were isolated from mLNs and pLNs of SPF mice. Subsequently, RNA-seq or ATAC-seq analyses were performed. DEGs and DARs were identified in mLN vs. pLN pairwise comparisons. Matrix layout for all intersections of putative TFs identified (bar graph left). Circles in the matrix indicate sets that are part of the intersection and the size of the intersection (all intersections with >= 2 are depicted) is indicated in the top bar graph. ATAC-seq, assay for transposase accessible chromatin sequencing; DAR, differentially accessible region; DEG, differentially expressed gene; mLN, mesenteric lymph node; pLN, skin-draining lymph node; SPF, specific pathogen-free; SC, stromal cell; TF, transcription factor.

**Supplementary Figure 2:**
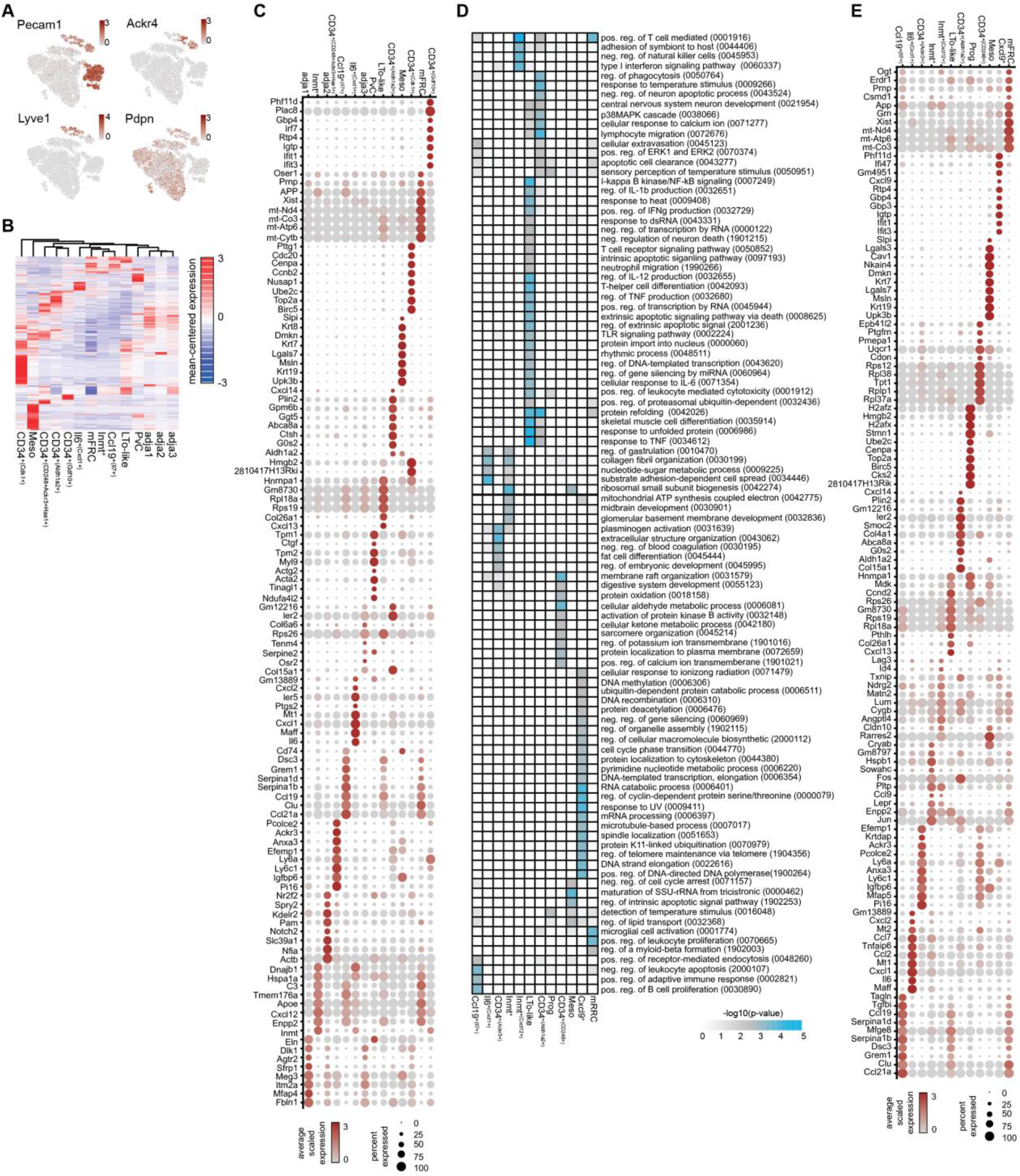
The LNSC compartment can be dissected into distinct cellular subsets using scRNA-seq. *Related to Figure 3.* CD45^−^CD24^−^ cells were isolated from mLNs of day 0, 10, 24, 56 and 300 old SPF-housed mice and subjected to scRNA-seq. **(A)** t-SNE plot of merged 15,925 endothelial and non-endothelial mLN SCs. Expression of indicated genes is superimposed on the t-SNE map. **(B)** Heatmap of the top 40 DEGs across all non-endothelial SC subsets including PvC and adjacent tissue cell subsets (adja). **(C)** Expression dot plot of top 8 DEGs, which were selected by foldchange, per subset comparing all remaining non-LEC and non-BEC mLN SCs. **(D)** GO analysis of biological processes was performed on DEGs per non-endothelial SC subsets. **(E)** Expression dotplot of top 10 DEGs, which were selected by foldchange, per subset comparing all non-endothelial SC subsets. DEG, differentially expressed gene; GO, gene ontology; Meso, mesothelial; mLN, mesenteric lymph node; Prog, progenitor; PvC, perivascular cells; SC, stromal cell; SPF, specific pathogen-free.

**Supplementary Figure 3:**
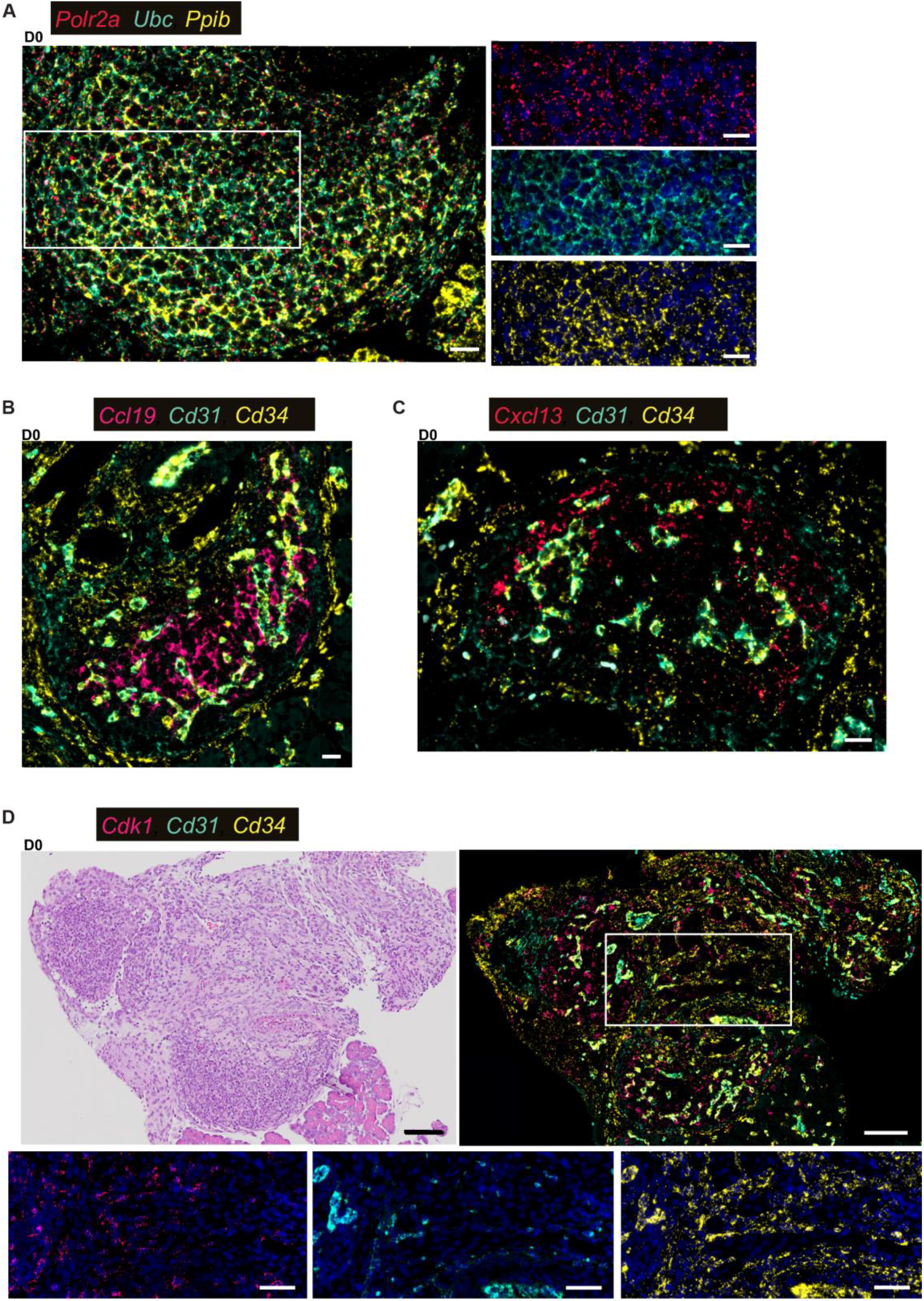
Expression of key marker genes in neonatal LN and adjacent tissues. *Related to Figure 3.* Sections (3 μm) of early postnatal mLN were stained with the indicated RNAscope probes and imaged by fluorescence microscopy. Nuclei were counter-stained with DAPI (blue). “White squares” indicate regions of interest. Representative tissue sections = 2-3. **(A)** Positive control probes for the genes *Polr2a*, *Ppib* and *Ubc* (scale bars = 20 μm). **(B)** Overview of mLN anlagen for RNA-probes specific for *Ccl19*, *Cd31* and *Cd34* (scale bar = 20 μm). **(C)** Overview of mLN anlagen for RNA-probes specific for *Cxcl13*, *Cd31* and *Cd34* (scale bar = 20 μm). **(D)** ***Upper left*** Haematoxylin-Eosin (HE) overview staining of D0 mLN embedded in surrounding tissue (scale bar = 100 μm). ***Upper right*** Overview of mLN anlagen for RNA-probes specific to *Cdk1, Cd31* and *Cd34* (scale bar = 100 μm). ***Lower*** Zoom-ins for RNA-probes specific to *Cdk1, Cd31* and *Cd34* (scale bar = 50 μm). HE, Haematoxylin-Eosin; mLN, mesenteric lymph node.

**Supplementary Fig. 4.**
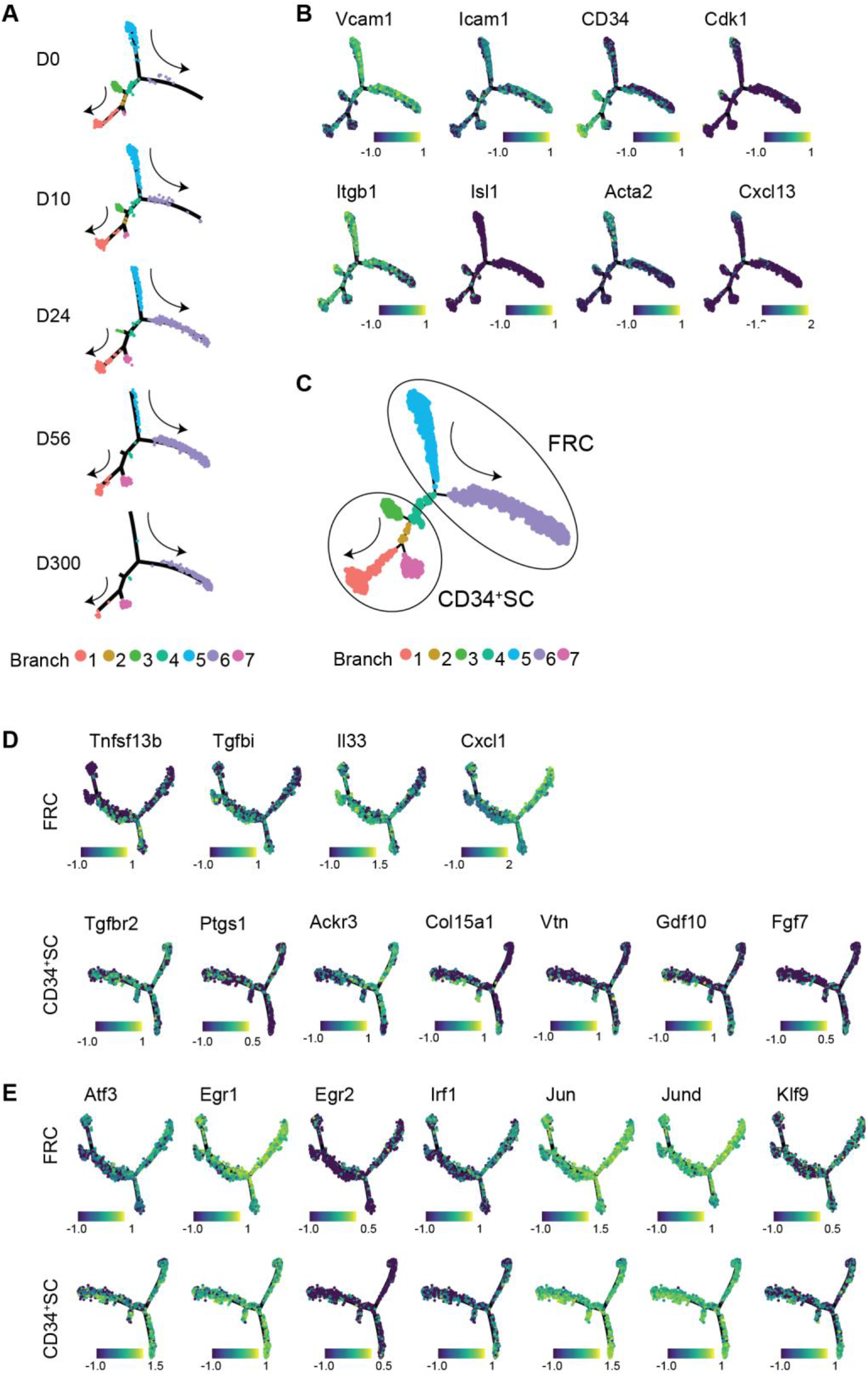
Bifurcation of LNSC development. *Related to Figure 4.* CD45^−^CD24^−^ cells were isolated from mLNs of day 0, 10, 24, 56 and 300 old SPF-housed mice and subjected to scRNA-seq. Target SCs were identified as non-LECs, non-BECs and non-PvCs. **(A)** Pseudotemporal trajectory of non-endothelial SCs plotted for the indicated age. Arrows indicate increasing pseudotime. **(B)** Gene expression on the pseudotime trajectory. **(C)** Applied electronic gating for re-embedding CD34^+^ SCs and FRCs. **(D)** Expression of selected DEGs along FRC and CD34^+^ SC. **(E)** Expression of common TFs for both FRCs and CD34^+^ SCs. DEGs, differentially expressed genes; FRC, reticular fibroblastic stromal cell; PvC, perivascular cell; SC, stromal cell; SPF, specific pathogen-free; TF, transcription factor.

## Methods

### Mice

CD90.1 mice (BALB/c) were bred and kept under SPF conditions in isolated ventilated cages at the Helmholtz Centre for Infection Research (Braunschweig, Germany). GF mice (BALB/c) were generated at Hannover Medical School (Hannover, Germany) by cesarean section and maintained either in plastic film isolators or in static micro-isolators (gnotocages) at Hannover Medical School or the Helmholtz Centre for Infection Research (Braunschweig, Germany). If not stated otherwise, water and food were supplied *ad libitum*. In all experiments, gender- and age-matched mice were used. All mice were housed and handled in accordance with good animal practice as defined by FELASA and the national animal welfare body GV-SOLAS.

### Antibodies

Fluorochrom-conjugated anti-human anti-CD24 (clone M1/69, APC, BioLegend Cat. #101814), anti-CD31 (clone 390, PE-Cy7, BioLegend Cat. #102418), anti-CD45 (clone 30-F11, APC, BioLegend Cat. #103112), anti-gp38 (clone 8.1.1, PE, BioLegend Cat. #127408) and Ter119 (clone Ly-76, APC, Biolegend Cat. #116212) were utilized in this study.

### RNAscope FISH

Triple hybridizations were carried out using the RNAscope Multiplex Fluorescent Detection Kit v2 (Advanced Cell Diagnostics, Cat. #320871 #300041 #323100 #310023 #310018) in combination with the corresponding 4-Plex Ancillary Kit (Advanced Cell Diagnostics, Cat #323120 #321831). The following target probes were used: Mm-Cdk1 (Cat. #476081, targeting bp 58-1159), Mm-Cxcl13 (Cat. #406311-C2, targeting bp 2-1143), Mm-Ccl19 (Cat. #432881-C2, targeting bp 5-712), Mm-CD34 (Cat. #319161-C3, targeting bp 383-1590) and Mm-CD31(Pecam1) (Cat. #316721-C4, targeting bp 915-1827). Sample preparation and stainings were carried out according to manufacturer’s instructions. In brief, mLNs from 0-1d old SPF-housed mice were dissected and fixed in 10% neutrally buffered formaldehyde for 16-32 h at RT, washed with 1x PBS (Gibco, Cat. #14190169), and dehydrated in a series of ethanol and xylene submersions before embedding in paraffin (Merck, Cat. #76242). Formalin-fixed paraffin-embedded (FFPE) tissue blocks were stored at 4°C. Sliced 3 μm tissue sections were continuously stored at 4°C until RNAscope stainings were performed. FFPE sections were baked at 60°C for 1 h, before being deparaffinized and dehydrated. Tissue sections were incubated with hydrogen peroxide for 10 min at RT before target retrieval was carried out in a steamer (Braun, Type 3216) for 15 min at >98°C. Protease treatment was performed with Protease Plus (Advanced Cell Diagnostics, Cat. #322380) for 20 min in a humidified hybridization chamber at 40°C. Subsequently, probes were allowed to hybridize to their targets for 2 h at 40°C in the hybridization chamber. During the following horse radish peroxidase (HRP) based amplification process, the tyramide signal amplification (TSA)-conjugated fluorophores Opal520, Opal570 and Opal650 (Perkin Elmer, Cat. #NEL80001KT) were used to visualize target probes. Tissue sections were counterstained with DAPI (Merck, Cat. #D9542) and mounted with ProLong Gold Antifade Mountant (Thermo Fisher Scientific, Cat. #P10144). Sections incubated with negative control probes (DapB) were stained in parallel and a mix of positive control probes (POLR2A, PPIB, UBC, Hprt) was utilized to confirm RNA integrity in each assessed tissue block. Images were acquired with an Olympus VS120 slide scanner fluorescence microscope using the VS-ASW-FL software (Olympus). Z-stacks were acquired at 40x or 20x magnification and extended focus imaging (EFI) was done at 20x magnification.

### Stromal cell isolation

For SC isolation, skin-draining pLNs (inguinal and axillary or popliteal) or mLNs (small intestinal and colon/caecum-draining) were resected and digested in RPMI 1640 medium (Gibco, Cat. #72400021) containing 0.2 mg/ml collagenase P (Roche, Cat. #11213865001), 0.15 U/ml dispase (Roche, Cat. #4942078001) and 0.2 mg/ml DNase I (Roche, Cat. #4536282001) as described previously.^34^ After digestion, cells were kept at 4°C in PBS containing 0.2 % BSA (Merck, Cat. #A2058) and 5 mM EDTA (Roth, Cat. #8043.1). CD45^−^ cells were enriched by autoMACS separation after magnetic labeling of CD45^+^ cells using anti-CD45-APC followed by anti-APC microbeads (Miltenyi Biotec, Cat. #130-090-855) or anti-CD45 Nanobeads (Biolegend, Cat. #480028). Subsequently, the CD45^−^ fraction was stained using fluorochrome-coupled antibodies and used to sort CD45^−^CD24^−^CD31^−^gp38^+^ FSCs (Aria II, 100 μm nozzle) and bulk CD45^−^CD24^−^ non-hematopoietic cells (Aria III, 70 μm nozzle) for RNA-seq, ATAC-seq and scRNA-seq.

### Transfection and lentiviral packing

Reverse lentiviral transfection was performed using Lipofectamine 2000 (Thermo Fisher, Cat. #11668027) following the manufacturer’s instruction. TF-bearing lentiviral vectors^47^ and lentiviral packaging plasmids pRSV-Rev (addgeneID 12253), pMDLg/pRRE (addgeneID 12251) and pCMV-VSV-G (addgeneID 8454) were supplemented at the ratio 1:1:1:1 and incubated for 30 min at the room temperature. Prior to transfection, HEK 293T cells (ATCC Cat. No. SD-3515) were washed with PBS (Thermo Fisher, Cat. #14190169) and dissociated using 0.25% Trypsin-EDTA (Life Technologies, Cat. #25200056). Cells were resuspended in DMEM (Gibco, 41966029) 10% FBS (Gibco, Cat. #10270106) and 1% penicillin-streptomycin (Life Technologies, Cat. #15140-122) and seeded in individual wells at confluence of 95% with the transfection mix. 12 h post-transfection, fresh medium was added to attached cells and the virus-containing supernatant collected after 48h, dead cells removed by centrifuging at 300 g for 10 min and supernatant stored at −80°C for up to 6 months.

### Lentiviral transduction

Murine C3H10T1/2 cells (ATCC Cat. No. CCL-226) were seeded 12 h prior to transduction at confluence of 10-20%. Transfection lentivirus-containing supernatant was mixed at 1:1 ratio with fresh medium and 10 μl/ml polybrene (Sigma, Cat. #TR-1003-G) and added to the plated adherent C3H10T1/2 cells. Cells were then centrifuged at 1300 g for 30 min at 37°C, incubated with the respective lentivirus for 24 h and fresh medium was added after 24 h. After 48 h, transduced cells were selected using 2 μl/ml of puromycin (Thermo Fisher, Cat. #A1113803) for 72 h. Subsequent to puromycin selection, medium was replaced and puromycin-resistant cells permitted to recover for 48 h. Then, medium containing 2 μl/ml doxycycline was added (Sigma, Cat. #D9891-1G) to induce TF expression. Medium with doxycycline was replenished every 48 h and doxocycline treatment maintained for 12 days. Direct-zol RNA kit (Zymo Research, Cat. #R2052) was used to extract RNA according to the manufacturer’s instruction.

### Library preparation WGBS

Genomic DNA was isolated from purified CD45^−^CD31^−^Ter119^−^gp38^+^ using the AllPrep DNA/RNA Micro Kit (Qiagen, Cat. #80284) according to the manufacturer’s instructions. Concentration and quality of the purified genomic DNA (gDNA) was determined by using NanoDrop (Thermo Fisher Scientific). Fragmentation of gDNA was carried out via Covaris S2 (Covaris), at duty cycle 10%, intensity 4 and 200 cycles per burst during 80 sec, to obtain fragments with an average length of 300 bp. The size of the fragments was verified with Agilent Technologies 2100 Bioanalyzer. DNA sequencing libraries were generated from fragmented gDNA using the TruSeq DNA Sample Prep Kit v2 (Illumina, Cat. #15026486) according to the manufacturer’s instructions and extending the workflow by adding one additional step: Subsequent to ligation of the adapter molecules to the DNA fragments, the sample was subjected to bisulfite conversion reaction using the EZ DNA Methylation Kit (Zymo Research, Cat. #D5001). The protocol for the True Seq DNA generation was then followed. The bisulfite-converted library was amplified by performing a PCR reaction (10 cycles, 98°C for 10 s, 63°C for 30 s, 72°C for 1 min) including the TruSeq primer mix and the KAPA Hifi Uracil+ Polymerase Master Mix (Kapa Biosystems, Cat. #KK2801). The PCR product was purified and size controlled by Agilent Technologies 2100 Bioanalyzer (High Sensitivity DNA Chip). The libraries were sequenced on an Illumina HiSeq2500 sequencer using the TruSeq SBS Kit v3-HS (200 cycles, paired end run) with an average of 2*10^8^ reads per replicate. The WGBS raw and processed data are available at NCBI GEO (GSE172526).

### Library preparation ATAC-seq

CD45^−^CD24^−^CD31^−^gp38^+^ cells were sorted by FACS into PBS containing 0.2% BSA. Cells were washed once with PBS before DNA transposition was performed with the Nextera DNA Library Prep Kit (Illumina, Cat. #FC-121-1031). Per sample, 25 μl TD, 2.5 μl TDE1 and 22 μl nuclease-free water were combined and placed at 37°C for 3 min before 0.5 μl of 1% Digitonin (Promega, Cat. #G9441) was added to the master mix. Samples were resuspended in the transposition reaction mix and incubated for 30 min at 37°C at 300 rpm. After transposition, DNA was purified with the MinElute PCR Purification kit (Qiagen, Cat. #28006) according to manufacturer’s instructions and eluted in 50 μl nuclease free water. Transposed DNA fragments were pre-amplified using 10 μl transposed DNA, 10 μl nuclease free water, 2.5 μl 25 μM custom Nextera PCR primer 1, 2.5 μl 25 μM custom Nextera PCR primer 2, 25 μl NEBNext High-Fidelity 2x PCR Master Mix (New England BioLabs, Cat. #M0541L) per reaction and amplified via a 6-cycle PCR program (1 cycle of 72°C for 5 min, 98°C for 30 s; 5 cycles of 98°C for 10 s, 63°C for 30 s, 72°C for 1 min). The forward primer was identical for all samples 5′-AATGATACG GCGACCACCGA GATCTACACTC GTCGGCAGCGT CAGATGTG-3′, whereas the reverse primer contained distinct barcodes (example underlined) used for demultiplexing 5′-CAAGCAGAAGA CGGCATACGAG AT*<u>TCGCCTTA</u>* GTCTCGTGGGC TCGGAGATGT-3′ ^62^. The appropriate amount of further amplification cycles was determined by qPCR using 5 μl of the pre-amplified product. Final amplification was carried out with 45 μl of previously PCR amplified DNA, 39.7 μl nuclease free water, 2.25 μl 25 μM customized Nextera PCR primer 1, 2.25 μl 25 μM customized Nextera PCR Primer 2, 0.81 μl 100x SYBR Green I and 45 μl NEBNext High-Fidelity 2x PCR Master Mix with the PCR program of 1 cycle of 98°C for 30 s; 8-10 cycles (depending on qPCR results) of 98°C for 10 s, 63°C for 30 s, 72°C for 1 min. PCR purification was carried out using the MinElute PCR Purification kit. Finally, size selection was performed with SPRIselect beads (Beckmann-Coulter, Cat. #B23317) with 1.2x for left-side and 0.55x for right-side selection according to manufacturer’s instructions. DNA quality, content and fragment size was assessed with Agilent Technologies 2100 Bioanalyzer profiles and Qubit measurements. Libraries were sequenced on an Illumina NovaSeq6000 sequencer using 50 bp single-end reads, and quality of sequenced libraries was verified with *FastQC*. The ATAC-seq raw and processed data are available at NCBI GEO (GSE172526).

### Library preparation RNA-seq

Total RNA was extracted from FACS-sorted CD45^−^CD24^−^CD31^−^gp38^+^ using the RNeasy Plus Micro Kit (Qiagen, Cat. #74034). cDNA was synthesized and amplified using template switching technology of the SMART-Seq v4 Ultra Low Input RNA Kit (Clontech Laboratories, Cat. #R400752), followed by purification using the Agencourt AMPure XP Kit (Beckman Coulter, Cat. #A63880). Library preparation was performed with Nextera XT DNA Library Prep Kit (Illumina). The Agilent Technologies 2100 Bioanalyzer was used to control quality and integrity of nucleic acids after each step. Deep sequencing was carried out on an Illumina HiSeq2500 sequencer using 50 bp single reads. Sequenced libraries were assessed for read quality using the FastQC tool. The RNA-seq raw and processed data are available at NCBI GEO (GSE172526).

### Library preparation scRNA-seq

Single CD45^−^CD24^−^ cells were sorted by FACS ARIA III (BD) and collected in PBS containing 0.04 % w/v BSA at a density of 400 cells/μl. Chromium™ Controller was used for partitioning single cells into nanoliter-scale Gel Bead-In-EMulsions (GEMs) and Single Cell 3’ reagent kit v2 for reverse transcription, cDNA amplification and library construction (10xGenomics, Cat. #120236). The detailed protocol was provided by 10xGenomics. SimpliAmp Thermal Cycler was used for amplification and incubation steps (Applied Biosystems). Libraries were quantified by Qubit™ 3.0 Fluometer (ThermoFisher) and quality checked using 2100 Bioanalyzer with High Sensitivity DNA kit (Agilent). Sequencing was performed in paired-end mode (2 x 75 cycles) on an Illumina NextSeq 500 sequencer to attain approximately 75,000 ± 25,000 reads per single cell. The scRNA-seq raw and processed data are available at NCBI GEO (GSE172526 and GSE106489 for D56 mLN-SPF^11^).

### Library preparation BRB-seq

3’end bulk mRNA cDNA library preparation and sequencing was performed following the BRB-seq strategy as previously described.^50^ In brief, 20 ng of total RNA isolated from TF overexpressions group, each represented with four to five replicates of two independent experiments, were reverse transcribed using SuperScriptTM II Reverse Transcriptase (Lifetech, Cat. Cat. #18064014) with individual barcoded oligo-dT primers, featuring a 12-nt-long sample barcode (IDT). Double-stranded cDNA was generated by second strand synthesis via the nick translation method. To that end, a mix containing 2 μl of RNAse H (NEB, Cat. #M0297S), 1 μl of E. coli DNA ligase (NEB, Cat. #M0205 L), 5 μl of E. coli DNA Polymerase (NEB, Cat. #M0209 L), 1 μl of 10mM dNTP (Thermo Fisher Scientific, Cat. #0181), 10 μl of 5x Second Strand Buffer (100 mM Tris, pH 6.9, [AppliChem, Cat. #A3452], 25 mM MgCl2 [Sigma, Cat. #M2670], 450 mM KCl [AppliChem, Cat. #A293], 0.8 mM β-NAD [Sigma, Cat. N1511], 60 mM (NH4)2SO4 [Fisher Scientific, Cat. #AC20587]), and 11 μl of water was added to 20 μl of ExoI-treated first-strand reaction on ice. The reaction was incubated at 16°C for 2.5 h. Full-length double-stranded cDNA was purified with 30 μl (0.6x) of AMPure XP magnetic beads (Beckman Coulter, Cat. #A63881) and eluted in 20 μl of water. cDNA concentration was measured using Qubit, and cDNA quality was assessed using a Fragment Analyzer (Agilent). cDNA was tagmented with in-house Tn5 ^63^, and libraries were purified using AMPure XP magnetic beads (0.6X). The resulting libraries were profiled with a High Sensitivity NGS Fragment Analysis Kit (Advanced Analytical, Cat. #DNF-474) and measured with the Qubit dsDNA HS Assay Kit (Invitrogen, Cat. #Q32851) prior to pooling and sequencing on an Illumina NextSeq 500 sequencer utilizing a BRB-seq custom primer and the High Output v2 kit (75 cycles) (Illumina, Cat. #FC-404-2005). The sequencing configuration is as follows: Read1 21cycles / index i7 8cycles / Read2 55c. The BRB-seq raw and processed data are available at NCBI GEO (GSE172526).

### WGBS analysis

The sequenced 2×100 bp paired-end libraries of bisulfite-treated reads were quality assessed and trimmed with the *FastQC* (version 0.11.1)^64^ and *Cutadapt* (version 1.4)^65^ tools. Trimmed libraries were aligned with the bisulfite short read mapping program *BSMAP* (version 2.4.3) (Xi & Li, 2009) versus the mouse reference genome (assembly: GRCm38), and the methylation status of each CpG site was called. Methylation profiles of CpGs with a minimum coverage of five mapped reads in at least two replicates of one condition and significant (based on t-statistics) change in their methylation status, served as input for the detection. Differentially methylated regions (DMRs) were identified using *bsmooth* with default settings.^28^

### ATAC-seq analysis

Sequencing reads were mapped to mouse genome (mm10) using *STAR* (version 2.5.3a)^66^ with parameters *--runMode alignReads --outSAMtype BAM SortedByCoordinate -- outFilterMultimapNmax 1* (assembly: GRCm38). Duplicates were removed using *picard MarkDuplicates* (https://broadinstitute.github.io/picard/). Peaks were called on de-duplicated bam-files using *macs2 callpeak* with the parameters *--broad -g mm -q 0.05* for each replicate.^67^ Heatmaps of fragment distribution around the TSS were computed using *computeMatrix* with the *reference-point -a 3000 -b 3000* and plotted using *plotHeatmap.* The regions identified via *macs2* were merged across all replicates into one set of regions, by combining peaks over-lapping with at least one base-pair and removing peaks that overlapped with blacklisted regions.^68^ Differential accessibility of raw ATAC-seq counts for each region/peak across all replicates of all samples were normalized across replicates with size factors computed with *DESeq2* (version 1.22).^69^ Pairwise comparisons were performed with *DESeq2* and differentially accessible regions (DARs) were called with an FDR adjusted p-value of less than 0.05 and a fold change (FC) of at least 2. Genomic features were identified via *getAnnotation* from *ChIPpeakANNO.*^70^ The cumulative FC of all DARs for one respective gene is represented as the mean of all the FC of all respective DARs. Transcription factor motif enrichment was computed using *homer* (version 4.9).^71^ Graphics were generated in R using *pheatmap* and *ggplot2*.

### RNA-seq analysis

Libraries were aligned versus the mouse reference genome assembly GRCm38 using the splice junction mapper Tophat2 v1.2.0 with default parameterization.^72^ Reads aligned to annotated genes were quantified with the HTSeq (version 0.12.4)^73^ and determined read counts served as input to DESeq2^69^ for pairwise detection and quantification of differential gene expression. RPKM (reads per kilobase of exon length per million mapped reads) values were computed for each library from the raw read counts. For scatterplots and heatmaps only genes with an annotated official *Gene Symbol* were included. Gene ontology (GO) analyses were performed using the R package *TopGo.*^74^ The R packages *pheatmap* and *ggplot2* were used to generate heatmaps or scatterplots, respectively.

### scRNA-seq analysis

Data were processed using Cell Ranger software (version 2.0.0) Count matrices were further processed with Seurat (version 2.3.3). All cells received an identifier which was used as common meta-data throughout the analysis including differentiation trajectories and dynamic gene regulatory networks (see below).

All cells with less than 1,000 or more than 4,600 detected genes per cell were filtered out. Moreover, cells with more than 4.5 % read mapping to mitochondrial genes were removed yielding 15,659 cells passing QC. After filtering, data were default normalized and the 2,000 most variable genes identified. The expression levels of these genes were scaled before performing PCA. The following covariates were regressed out: number of UMIs, percent of mitochondrial reads, percent of ribosomal reads and scores for the proliferation S.Score and G2M.Score computed with *CellCycleScoring()*. t-SNE dimensionality reduction was performed using the first 12 dimensions of the PCA and resolution set to 1.1. Only clusters (non-endothelial SC) with normalized expression for *Pecam* < 1 were used for the further analysis amounting to 9,323 cells which were re-embedded as described above (resolution = 1.0). The perivascular mesenchymal cell (PvMC) cluster and clusters classified as adjacent cells were excluded and the remainder cells, numbering 5,658 mesenchymal cells were re-embedded as described above (resolution = 1.0). For subsetting the *Prog.* subset, 259 cells were re-embedded as described above (resolution = 0.5, dimsuse = 10). GO analysis was performed for differentially upregulated genes per cluster using *TopGO.*^74^

Differentiation trajectories were analyzed using *Monocle* (version 2.18.0).^42^ Unsupervised ordering was performed on the 5,658 mesenchymal cells using *Monocle2’s DDRTree* algorithm based on genes with a mean expression >0.1. The trajectory containing all mesenchymal cells, was split based on marker gene expression *Vcam1* and *Cd34* and the annotation of cells belonging to the Prog^Cxcl13+^ or Prog^CD34+^ subset to distinct terminal branches. Two separate unsupervised orderings were performed as described above and are referred to as CD34^+^ SC or FRC trajectory.

Gene regulatory networks were inferred with the *dynGENIE3* algorithm,^46^ where the input expression values were based on ordering the genes according to physiological age per branch for each of the two trajectories being CD34^+^ SC or FRC. The list of candidate TFs was derived from TFBS from ATAC-seq profiling or DMRs within the proximity of TSSs. For network visualization with Cytoscape (version 3.6.0),^75^ only the top 500 links ranked by weight assigned by *dynGENIE3* were used. Node centrality and betweenness were calculated with the degree and betweenness functions from the igraph (version 1.2.2) package.

### BRB-seq analysis

After sequencing and standard Illumina library demultiplexing, the fastq-files were aligned to the mouse reference genome mm10 (GRCm38) using STAR (version 2.7.3a), excluding multiple mapped reads. Resulting BAM files were demultiplexed per sample using BRB-seqTools (version 1.4, https://github.com/DeplanckeLab/BRB-seqTools) and read-count matrices generated using HTSeq (version 0.12.4). Raw read counts were converted to transcripts per kilobase of exon per million reads values. Protein-coding genes with at least 5 reads in at least two replicates were included in the analysis. The calculated read counts were further processed with *DESeq2* for quantification of differential gene expression.^69^ Genes were considered as differentially expressed at fold change > 2.0 and the FDR adjusted p-value of < 0.05.

### Statistical analysis

For all scripts written in R, we used version 3.4.1 unless otherwise noted.

## Data Availability

ATAC-seq, WGBS, RNA-seq, scRNA-seq and BRB-seq raw and processed data generated during this study are available at NCBI GEO (GSE172526).

## Code Availability

Code used for this study will be made available upon request.

## Acknowledgements

We thank the Cell Sorting facility (HZI) for cell sorting, the Genome Analytics facility (HZI) for RNA-seq, scRNA-seq, WGBS and ATAC-seq sequencing, and the Genome Analytics facility (EPFL-SV) for BRB-seq sequencing. We thank the Department of Experimental Immunology and especially Dr. Lothar Gröbe and Maria Höxter for their technical assistance. This work was supported by the Hannover Biomedical Research School (HBRS), the Center for Infection Biology (ZIB) from Hannover Medical School, as well as the Deutsche Forschungsgemeinschaft (DFG, German Research Foundation) under Germany’s Excellence Strategy – EXC 2155 “RESIST” – Project ID 39087428, SPP1656 (Ho2236/9-2), and PE 2840/1-1). P.A. was supported by the Interdisciplinary Center for Clinical Research (IZKF) of the University Hospital of Wuerzburg (Project Z-6). The Helmholtz Institute for RNA-based Infection Research (HIRI) supported this work with a seed grant through funds from the Bavarian Ministry of Economic Affairs and Media, Energy and Technology (Grant allocation nos. 0703/68674/5/2017 and 0703/89374/3/2017).

## Author contributions

J.P., C.W., M.Bi., M.Z., W.C., S.F., M.E., J.R. and P.A. performed experiments and interpreted data. J.P., M.L., V.G., M.Be., C.W. and E.V. performed bioinformatics analysis. B.D., K.S., A.-E.S. and W.C. provided expertise and feedback. J.P., C.W., and J.H. designed research, interpreted data, and wrote the manuscript.

